# TRIM5α restricts flavivirus replication by targeting the viral protease for proteasomal degradation

**DOI:** 10.1101/605345

**Authors:** Abhilash I. Chiramel, Nicholas R. Meyerson, Kristin L. McNally, Rebecca M. Broeckel, Vanessa R. Montoya, Omayra Méndez-Solís, Shelly J. Robertson, Gail L. Sturdevant, Kirk J. Lubick, Vinod Nair, Brian H. Youseff, Robin M. Ireland, Catharine M. Bosio, Kyusik Kim, Jeremy Luban, Vanessa M. Hirsch, R. Travis Taylor, Fadila Bouamr, Sara L. Sawyer, Sonja M. Best

## Abstract

Tripartite motif-containing protein 5α (TRIM5α) functions as a cellular antiviral restriction factor with exquisite specificity towards the capsid lattices of retroviruses. The relative avidity of TRIM5α binding to retrovirus capsids directly impacts primate species susceptibility to infection, but the antiviral role of TRIM5α is thought limited to retroviruses. In contrast to this current understanding, here we show that both human and rhesus TRIM5α possess potent antiviral function against specific flaviviruses through interaction with the viral protease (NS2B/3) to inhibit virus replication. Importantly, TRIM5α was essential for the antiviral function of IFN-I against sensitive flaviviruses in human cells. However, TRIM5α was ineffective against mosquito-borne flaviviruses (yellow fever, dengue, and Zika viruses) that establish transmission cycles in humans following emergence from non-human primates. Thus, TRIM5α is revealed to possess remarkable plasticity in recognition of diverse virus families, with potential to influence human susceptibility to emerging flaviviruses of global concern.

## Main

Flaviviruses (family *Flaviviridae*) include 53 recognized virus species of which 40 are known to cause disease in humans, with over 40% of the world’s population at risk of flavivirus infection annually ^1^. These viruses have high potential for emergence into human populations as witnessed historically through global emergence of dengue virus (DENV), West Nile virus (WNV), and Zika virus (ZIKV). Additional (re)emerging viruses of considerable medical importance include yellow fever virus (YFV), Japanese encephalitis virus (JEV) and members of the tick-borne encephalitis virus (TBEV) serogroup. Flaviviruses share in common a positive-sense single-stranded RNA (ssRNA) genome encoding a single polyprotein that is cleaved by host cell signalases ^2^ and the viral protease to generate three structural (capsid [C], pre-membrane [M] and envelope [E]) and seven nonstructural (NS) proteins (NS1, NS2A, NS2B, NS3, NS4A, NS4B and NS5) ^3^. Two of the nonstructural proteins have enzymatic activity; the NS3 protein encodes the viral RNA helicase and together with its co-factor NS2B (NS2B/3) functions as the viral protease, whereas NS5 possesses both methyltransferase (MTase) and RNA-dependent RNA polymerase (RdRP) activities.

The repeated emergence of flaviviruses as human pathogens is in part due to the fact that they are arthropod-borne, transmitted by mosquitoes and ticks. In addition, the zoonotic reservoir species’ supporting virus replication in nature are highly diverse. For example, small mammals, particularly rodents, are thought critical for maintenance of transmission cycles of TBEV and related viruses. In contrast, WNV utilizes birds, whereas DENV, ZIKV and YFV evolved in non-human primates before at least DENV and ZIKV established urban transmission cycles maintained exclusively through human infections ^4^. The ability of a virus to avoid or evade host antiviral responses is essential to establish replication and transmission ^5^. However, it is not fully understood how evolution in different reservoir hosts to avoid innate immunity has shaped replication and pathogenesis of different flaviviruses following infection of humans. Host-specific interactions with the interferon (IFN) response have been demonstrated for DENV and ZIKV that can only antagonize IFN-dependent signaling in the context of primate hosts ^6^. However, the IFN-stimulated genes (IGSs) that might also contribute to host-specific restriction of flaviviruses are not well characterized.

Tripartite motif-containing proteins (TRIMs) are strong candidates for mediating host-specific restriction of virus replication in the context of an IFN response. Approximately 100 tripartite TRIMs exist in the human genome ^7^, many of which are ISGs with functions as direct antiviral restriction factors or as modulators of the cellular response to infection ^8^. The most characterized primate TRIM is TRIM5α, which functions as a cellular antiviral restriction factor with exquisite specificity, thought to restrict only retroviruses through complex interactions with the capsid lattice structure that accelerates uncoating of the viral nucleic acid and also blocks reverse transcription ^9–11^. The significant influence of TRIM5α is exemplified by the observations that its antiviral activity drives lentivirus evolution ^9^ and limits cross-primate species transmission ^12^. Importantly, the relative ability of TRIM5α to bind retrovirus capsid lattices directly impacts primate species susceptibility to infection. For example, TRIM5α from Old World monkeys such as rhesus macaques (rhTRIM5α) exerts potent antiviral activity against HIV-1 to confer host resistance. In contrast, human TRIM5α (hTRIM5α) only weakly interacts with HIV-1 capsid lattices and this reduced efficacy may promote HIV-1 transmission and disease progression ^13^. The antiviral specificity of TRIM5α has evolved rapidly in the past 30 million years of primate evolution, with particularly strong signatures of positive selection over the last 4-5 million years ^14,15^. Evolutionary studies support the conclusion that TRIM5α positive selection throughout primate evolution is driven at the interaction interface between TRIM5α and retrovirus capsids, and thus reinforce the paradigm that the antiviral activity of TRIM5α and its role in host resistance is specific to the retroviruses ^16^.

Given the extensive evolution of multiple medically important flaviviruses with primate species, we examined the antiviral capacity of both rhTRIM5α and hTRIM5α towards the vector-borne flaviviruses. Surprisingly, both rhTRIM5α and hTRIM5α possessed potent antiviral function against specific flaviviruses within the TBEV serogroup, but not towards mosquito-borne flaviviruses. The antiviral activity of TRIM5α was mediated through interactions with the viral protease NS2B/3 at sites of virus replication, and association of TRIM5α with NS2B/3 from a sensitive virus resulted in proteasomal degradation of the viral protein. Importantly, human TRIM5α contributed significantly to the antiviral effects of type I IFN against sensitive tick-borne viruses. However, TRIM5α was ineffective against important mosquito-borne flaviviruses including YFV, DENV, and ZIKV. Thus, this work reveals an unexpected role for primate TRIM5α as an anti-flavivirus restriction factor that may influence human susceptibility to infection.

## RESULTS

### TRIM5α is a functional restriction factor for flaviviruses

The association of various mosquito-borne flaviviruses with primates prompted us to test whether ectopic expression of TRIM5α might have anti-flavivirus activity. HEK293 cells were engineered to stably express various TRIM5α proteins (Supplementary Fig. 1a). Expression of rhesus macaque (rh) TRIM5α restricted infection of vesicular stomatitis virus glycoprotein (VSV-G) pseudotyped HIV-1 in 293 cells, demonstrating that these cells are appropriate to observe TRIM5-mediated restriction (Supplementary Fig. 1b). Compared to empty vector control cells, expression of either human (h) TRIM5α or rhTRIM5α restricted replication of related viruses in the TBEV serogroup, including TBEV (strain Sofjin), Kyasanur Forest disease virus (KFDV) and Langat virus (LGTV; an attenuated member of the TBEV serocomplex), but not WNV (strain NY99), DENV (strain NGC, serotype 2), ZIKV (strain 2013 French Polynesia) or YFV (strain 17D) (Fig. 1a). TRIM5α did not affect replication of Powassan virus (POWV; strain LB) despite this virus also belonging to the TBEV serogroup. The impact of hTRIM5α or rhTRIM5α on replication of sensitive flaviviruses was significant, reducing production of infectious virus by up to 1000-fold during the exponential phase of virus growth. hTRIM5α was functional but less efficient, imposing a 90% reduction but this may be attributable to the lower expression of hTRIM5α compared to rhTRIM5α (Supplementary Fig. 1a). Therefore, we also used CrFK cells stably expressing hTRIM5α-HA as a cell model historically used to examine retrovirus restriction as they lack intrinsic TRIM5α expression ^17^. Expression of hTRIM5α suppressed replication of both TBEV (Supplementary Fig. 1c) and LGTV (data not shown), but not WNV (Supplementary Fig. 1d). In HEK293 cells that support more optimal flavivirus growth, restriction was observable up to a starting multiplicity of infection (MOI) of 10 (Supplementary Fig. 1e), but replication of TRIM5α-sensitive viruses eventually overcame restriction which is consistent with viral saturation of antiviral restriction factors ^18^ (Fig. 1a). A related human TRIM with anti-retrovirus function, TRIM22 ^19^, did not impact replication of TBEV, KFDV or LGTV, demonstrating a specific role for TRIM5α in flavivirus restriction (Fig. 1a). Suppressed replication of KFDV was also observed at the level of protein expression, with reduced accumulation of NS3 in cells expressing hTRIM5α-HA or rhTRIM5α-HA compared to the empty vector controls (Fig. 1b). Expression of the envelope protein (E) of sensitive viruses was also reduced when examined by flow cytometry (Fig. 1d). However, no reduction in either NS3 by Western blot or E expression by flow cytometry was observed following POWV infection, supporting flavivirus species-specific restriction by TRIM5α (Fig. 1c, d).

**Fig. 1.**
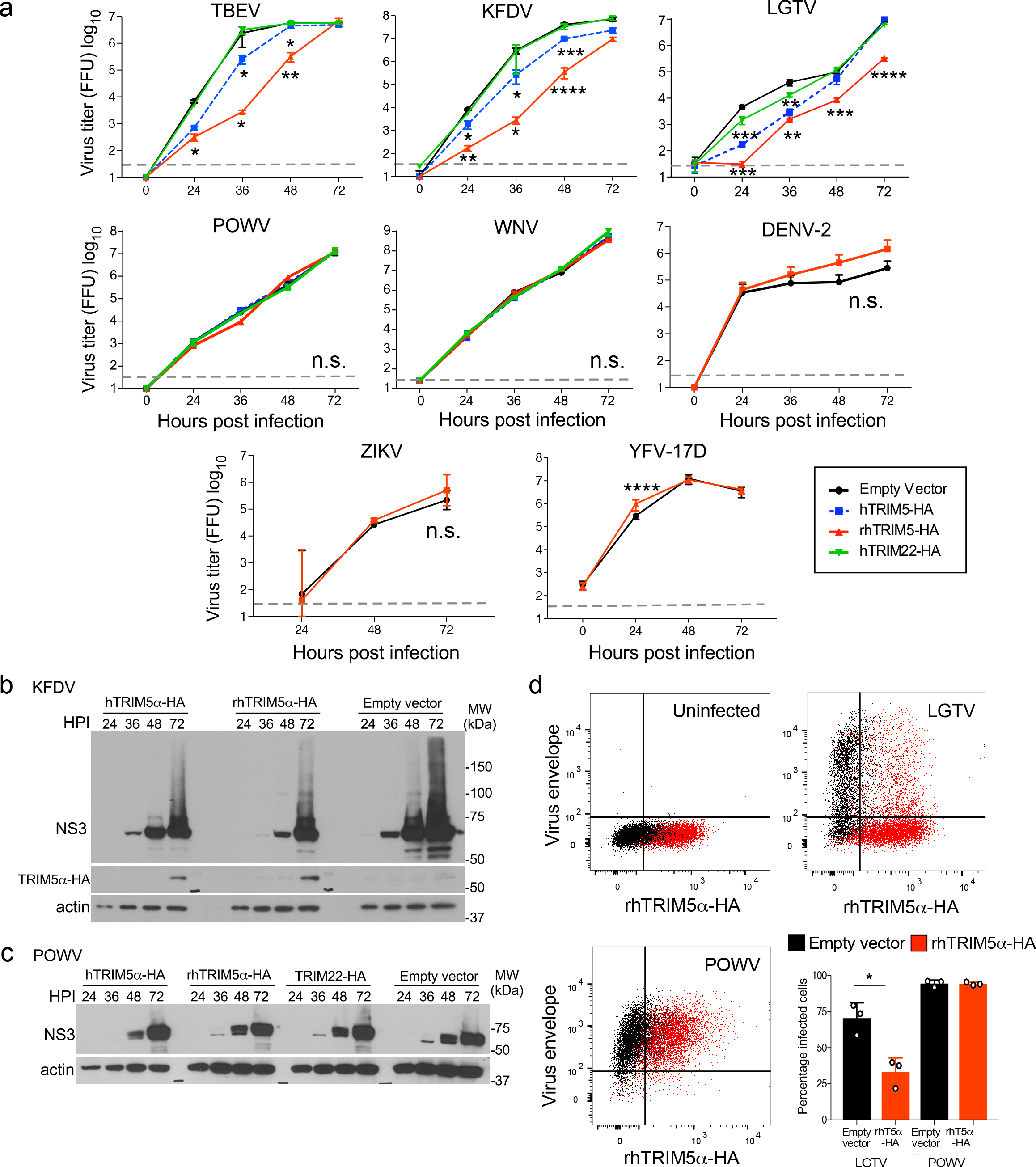
Stable expression of TRIM5α in HEK293 cells restricts replication of specific flaviviruses. **a**, HEK293 cells stably overexpressing human (h) or rhesus (rh) TRIM5α-HA, hTRIM22-HA or empty vector (control) were infected with tick-borne encephalitis virus (TBEV), Kyasanur Forest disease virus (KFDV), Langat virus (LGTV), Powassan virus (POWV), West Nile virus (WNV), dengue virus (DENV-2), Zika virus (ZIKV) or yellow fever virus (YFV) with a multiplicity of infection (MOI) of 0.001 (except YFV at MOI 0.1). Infectious virus release was determined in supernatants by plaque assay. All data are from three independent experiments performed in triplicate (mean ± s.d., *P < 0.05, **P < 0.01, ***P < 0.001, **** P < 0.0001, n.s. not significant). Grey dotted line indicates limit of detection. **b-c**, NS3 protein levels in stable HEK293 cells infected with **b**, KFDV, or **c**, POWV. **d**, Dot plots depicting an overlay of E protein in empty vector (black) or rhTRIM5α-HA cells (red) infected with LGTV or POWV measured by flow cytometry. The percentage of cells infected as measured by E protein staining is quantified in the bar graphs.

To determine if human TRIM5α is a functional restriction factor, *TRIM5* mRNA was depleted by RNA interference (RNAi) in A549 cells using lentivirus-delivered short hairpin RNA (shRNA), or *TRIM5* was knocked out using CRISPR/Cas9 in Hap1 cells. Cells were left untreated, or treated with IFNβ for 6 h prior to infection to upregulate *TRIM5* expression and induce an antiviral state. Reduced *TRIM5* expression did not affect the responsiveness of cells to IFNβ as measured by upregulation of canonical IFN-stimulated genes (ISGs), *RSAD2* (viperin) and *CXCL10* (Fig. 2a). However, depletion of *TRIM5* partially relieved the antiviral effect of IFNβ on LGTV (Fig. 2a). Transfection of A549 cells with an independent siRNA sequence targeted towards hTRIM5α also increased replication of LGTV but not YFV (Fig. 2b). Furthermore, deletion of TRIM5 using CRISPR/Cas9 in Hap1 cells (Supplementary Fig. 1f,g) rescued ~2 log_10_ LGTV replication in the presence of IFNβ (Fig. 2c). Virus replication was also increased for TBEV, but not POWV, WNV, ZIKV, DENV or YFV (Fig. 2c). Together, these results identify TRIM5α as a restriction factor for specific species of flaviviruses, and demonstrate that TRIM5α is an effector of the human type I IFN response to these viruses.

**Fig. 2.**
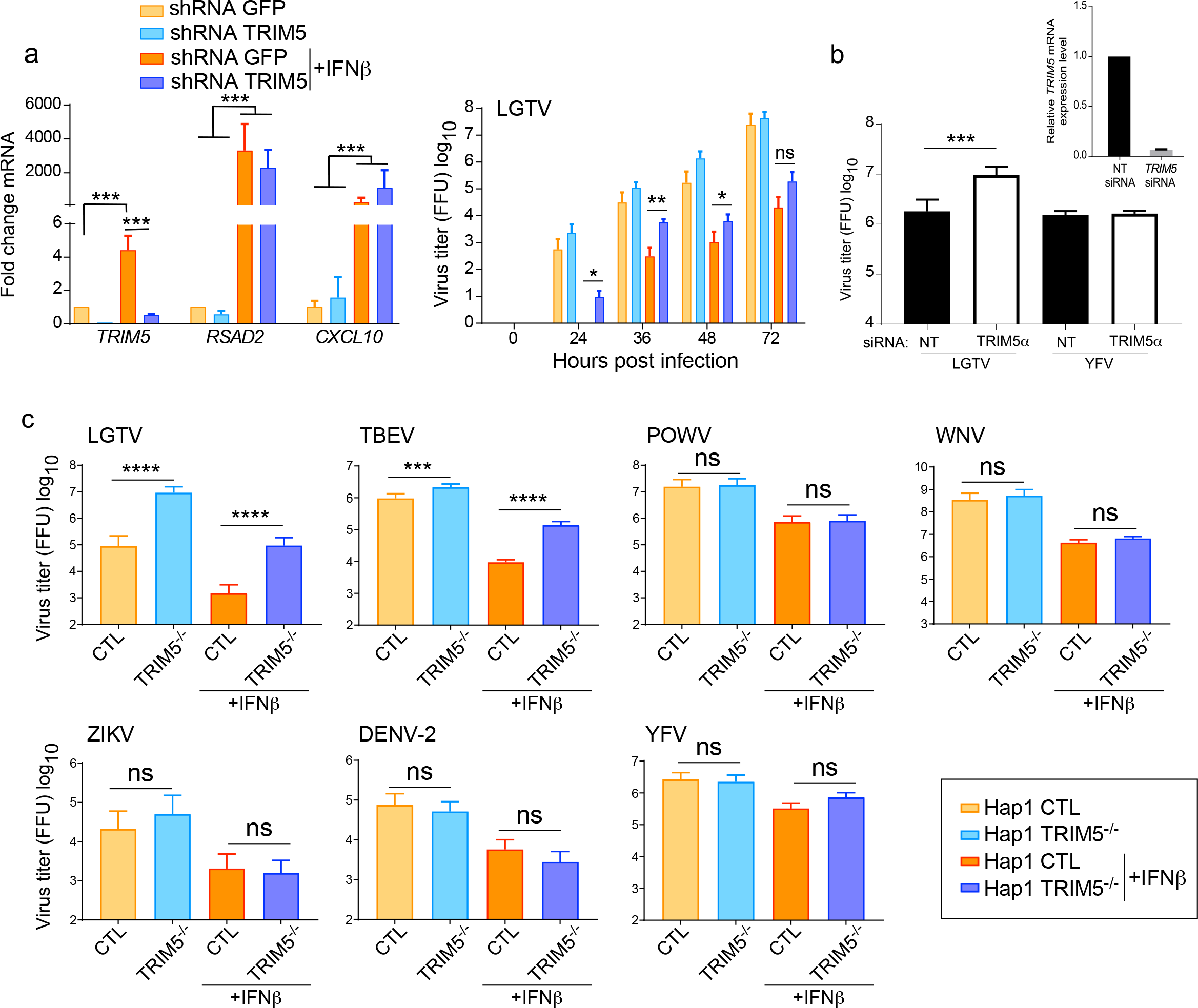
Endogenous human TRIM5 is an ISG required for the antiviral effects of IFNβ against TBEV and LGTV. **a**, Left panel: qRT-PCR for *TRIM5*, *RSAD2* or *CXCL10* mRNA isolated from A549 cells following transduction with lentiviruses expressing short hairpin RNAs (shRNAs) for GFP (control) or *TRIM5*, and untreated or treated with IFNβ (IFN) at 1000 U/ml for 6 h. Right panel: LGTV titers in A549 cells that were left untreated or pre-treated with IFNβ for 6 h and infected at MOI-0.001. Supernatants were collected at the indicated times and titrated by plaque assay. All data are from three independent experiments performed in triplicate (mean ± s.d., *P < 0.05, **P < 0.01 by Mann-Whitney; ns, not significant). **b**, A549 cells were transfected with siRNAs specific for *TRIM5* or a non-targeting (NT) control. Cells were infected with LGTV or YFV at 48 h post-transfection (MOI 0.001), and supernatants harvested for virus titration 48 h later. Data are from 3 independent experiments (mean ± s.d.; ***P < 0.001 by Mann-Whitney). Inset shows the relative *TRIM5* mRNA expression measured by qRT-PCR in A549 cells. **c**, Replication of LGTV, TBEV, POWV, WNV, ZIKV, DENV-2 and YFV (all infected at MOI 0.1) in Hap1 cells with *TRIM5* gene disruption by CRISPR/Cas9. Hap1 cells were left untreated or pretreated for 6 h with IFNβ. Data are from 2-3 independent experiments performed in triplicate (mean ± s.d., ***P < 0.001, **** P < 0.0001 by one-way ANOVA with Tukey’s multiple comparisons post-test; ns, not significant).

### TRIM5α expression restricts viral RNA replication

To determine which step in the flavivirus life cycle was restricted by TRIM5α, LGTV replication was examined in rhTRIM5α-HA HEK293 cells. At 48 hours post infection (hpi) supernatants and cell lysates were subjected to three cycles of freeze-thaw lysis to compare levels of intra- and extra-cellular virus. In the presence of rhTRIM5α, no change in the ratio (~1:10) of intracellular:extracellular infectious virus was observed (Fig. 3a) although intracellular accumulation of positive-sense (genomic) viral RNA was reduced by approximately 50-fold (Fig. 3b). Viral entry was not affected as differences in positive-sense RNA were not apparent after virus entry until at least 8-12 hpi when flavivirus RNA replication is initiated ^3,20^(Fig. 3c). Thus, TRIM5α imposes a block in virus replication at or preceding RNA replication without affecting virus entry or release. In flavivirus-infected cells, cellular localization of dsRNA is an obligate marker of sites of replication, and most perinuclear foci containing NS3 (the viral protease and RNA helicase) also colocalize with dsRNA, suggesting these perinuclear foci are sites of active replication ^21^ (Supplementary Fig. 2a). In infected cells, small aggregates of rhTRIM5α often termed cytoplasmic bodies ^22^ colocalized with NS3 and dsRNA suggesting that TRIM5α is recruited to replication complexes (Fig. 3d). RhTRIM5α aggregates also colocalized with NS5 (the viral RdRP) but only at perinuclear sites likely together with NS3 at the ER (Fig. 3e). Recruitment of human TRIM5α to sites of NS3 expression was also observed in LGTV-infected cells (Supplementary Fig. 2b). Areas of colocalization were observable between TRIM5α and dsRNA in the context of DENV or ZIKV, but unlike LGTV, infection did not induce strong aggregation of TRIM5α (Supplementary Fig. 3a,b). Next, we validated the association of either hTRIM5α or rhTRIM5α with NS3 by immunoprecipitation (IP) in LGTV-infected cells (Fig. 3f). Despite low levels of viral protein associated with restriction, IP of NS3 from infected cells resulted in co-precipitation with either hTRIM5α or rhTRIM5α (Fig. 3f). As expected, NS5 also co-precipitated with NS3 in infected cells which supports the IFA data and suggests that TRIM5α interactions with NS3 occur at sites of virus replication where NS3 and NS5 interact. Consistent with lack of TRIM5α aggregation at sites of dsRNA staining (Supplementary Fig. 2c, d), IP of NS3 from WNV-infected cells did not result in co-precipitation of rhTRIM5α (Supplementary Fig. 2e). Thus, TRIM5α localizes to viral replication complexes and suppresses RNA replication in a flavivirus-specific manner.

**Fig. 3.**
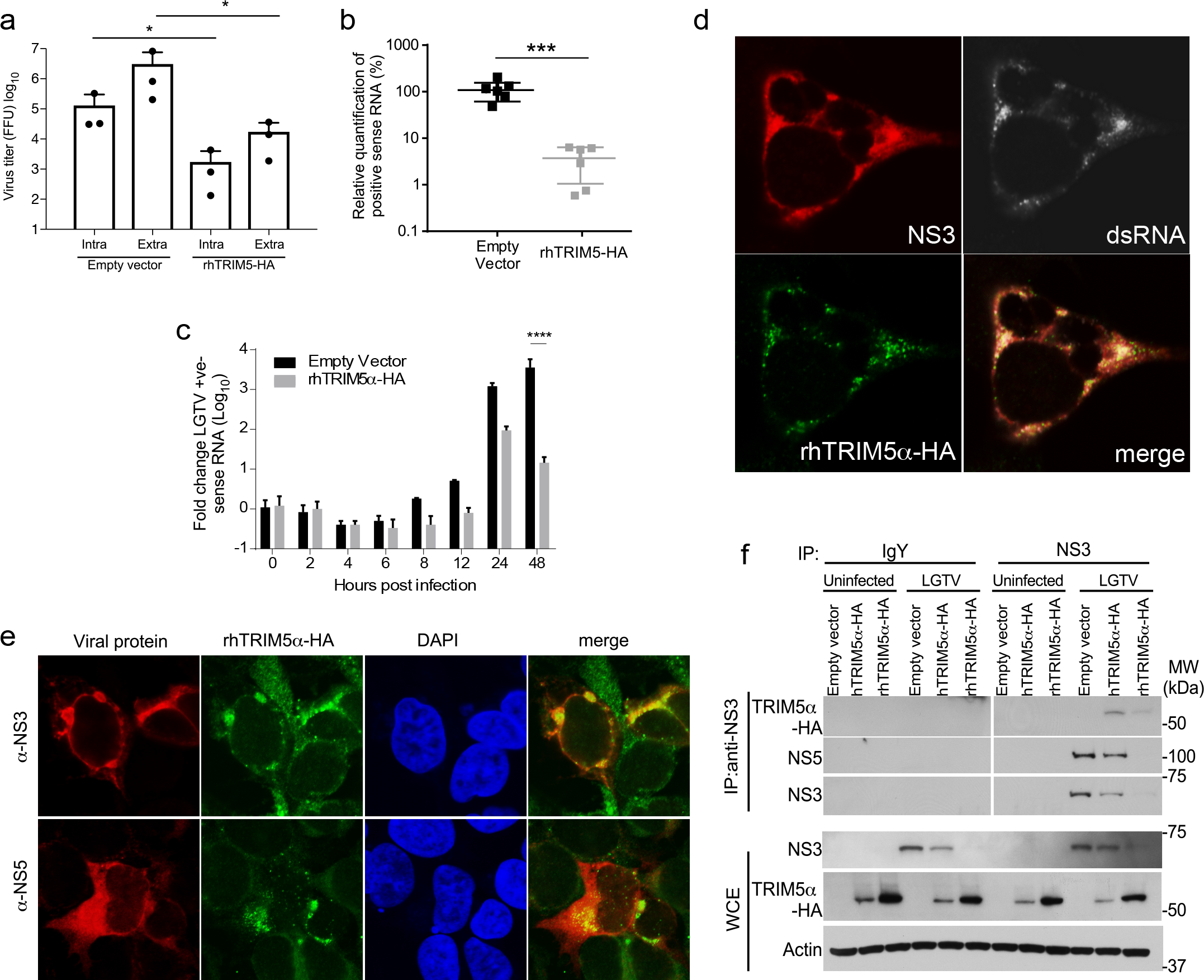
TRIM5α restricts flavivirus RNA replication and co-precipitates with the viral protease NS2B/3. **a**, HEK293 cells with stable expression of rhTRIM5α-HA or the empty vector (control) were infected with LGTV (MOI of 0.01). Infectious virus in cell supernatants or intracellular virus was quantified by plaque assay at 48 hpi. **b**, Accumulation of LGTV positive-sense viral RNA in cells infected in part A. was determined at 48hpi by qRT-PCR (mean ± s.d., *P<0.05; ***P < 0.001). **c**, Changes in genomic RNA over time following binding of LGTV to control and rhTRIM5α-HA-expressing HEK293 cells at 4°C and three washes with DPBS (mean ± s.d., ****P<0.0001 2-way ANOVA with Sidak’s posttest). **d**, Colocalization of NS3 (red), dsRNA (greyscale) and rhTRIM5α (green) in HEK293 rhTRIM5α-HA LGTV-infected cells at 24 hpi by IFA (MOI of 5). **e**, Colocalization of NS3 (red) or NS5 (red), and rhTRIM5α (green) in HEK293 rhTRIM5α-HA LGTV-infected cells at 24 hpi by IFA. Nuclei are counterstained with DAPI (blue) (MOI of 5). **f**, Interactions between rhTRIM5α or hTRIM5α with NS3 at 48 hpi with LGTV shown by immunoprecipitation (IP) of NS3 from infected HEK293 cells. WCE, whole cell extract.

### TRIM5α targets the flavivirus protease for proteasomal degradation

To examine interactions with NS3 and NS5 separately, stable rhTRIMα-HA cells were transfected with plasmids encoding LGTV NS2B/3 or NS5. NS2B was included as it forms an integral structural component of the NS3 protease active site, and transmembrane domains within NS2B target NS3 to ER membranes, with NS2B/3 being an important antiviral drug target ^23^. LGTV NS5 showed some co-localization with rhTRIM5α (Fig. 4a) and caused low levels of TRIM5α aggregation (Fig. 4b) but did not co-precipitate (Supplementary Fig. 4a). However, LGTV NS2B/3 expression caused rhTRIM5α to aggregate into discrete cytoplasmic bodies (Fig. 4a,b and Supplementary Fig. 4b) and co-localize reminiscent of that observed following virus infection, and LGTV NS2B/3 strongly associated with rhTRIM5α by co-precipitation (Fig. 4g). In addition, expression levels of NS2B/3 were reduced in cells expressing rhTRIM5α compared to the control cell line, whereas NS5 levels were not strongly affected (Fig. 4c). To further explore this observation, a constant level of LGTV NS2B/3 was expressed with increasing amounts of either rhTRIM5α or hTRIM5α by transfection of expression plasmids. In either case, expression of both unprocessed NS2B/3 and NS3 generated through autonomous cleavage was reduced in a dose-dependent fashion (Fig. 4d,e), although this effect was quickly saturated. Again, expression of LGTV NS5 was not affected by rhTRIM5α expression (Fig. 4f).

**Fig. 4.**
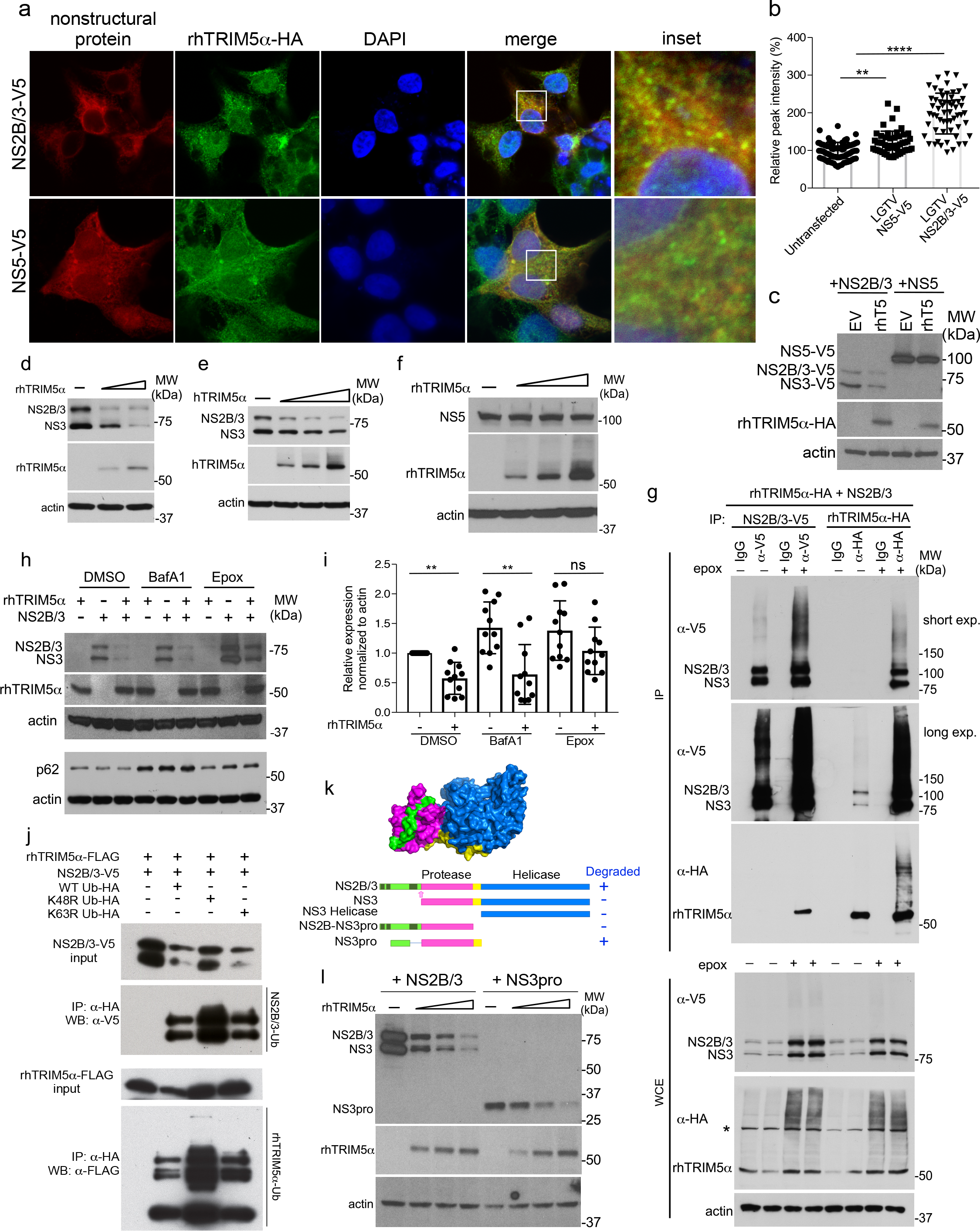
Binding of the flavivirus protease by rhTRIM5α is conformation dependent and results in proteasome-dependent degradation of NS2B/3. **a**, Stable HEK293 rhTRIM5α-HA (green) cells were transfected with plasmids coding for either NS2B/3-V5 or NS5-V5 (red) from LGTV and imaged by confocal microscopy. **b**, Relative intensity of TRIM5 aggregates were measured along vectors drawn in cells expressing LGTV NS2B/3 or NS5, with example vectors shown in Figure S3B. **c**, Western blot of LGTV NS2B/3-V5 or NS5-V5 in stable rhTRIM5α-HA or control HEK293 cells. **d-f.** Western blot analysis of HEK293 cells transfected with **d**, increasing amounts of rhTRIM5α-HA and constant amounts of LGTV NS2B/3-V5, **e**, increasing amounts of hTRIM5α-HA and constant amounts of LGTV NS2B/3-V5, **f**, increasing amounts of rhTRIM5α-HA and constant amounts of LGTV NS5-V5. **g**, Reciprocal co-IP of rhTRIM5α-HA and LGTV NS2B/3-V5 following cotransfection and 4 h treatment with epoxomicin (200 nM). The asterisk indicates a non-specific band. **h.** Western blot of LGTV NS2B/3-V5, rhTRIM5α-HA and endogenous p62 in HEK293 cells following 4 h treatment with DMSO (vehicle), Baf-A1 (200 nM) or epoxomicin (200 nM). **i**, Quantification of LGTV NS2B/3 expression with or without rhTRIM5α and treated with Baf-A1 or epoxomicin from 11 individual experiments. **j**, LGTV NS2B/3-V5 and rhTRIM5α-FLAG were co-expressed with ubiquitin (Ub)-HA WT or K48R or K63R mutants in HEK293 cells. Target proteins were immunoprecipitated using anti-V5 or anti-FLAG antibodies, and blots probed with anti-HA to examine Ub conjugation. **k**, Domain structure of flavivirus NS2B/3 (PDB: 2vbc) and schematic representation of truncation mutants. **l**, Western blot analysis of HEK293 cells transfected with increasing amounts of rhTRIM5α-HA and constant amounts of LGTV NS3pro. Lysates were probed specifically for HA, V5 and b-actin.

In the context of HIV-1, TRIM5α utilizes the proteasome (MG132 sensitive) for capsid disruption but not for restriction ^24^, and may also use lysosomes following autophagy (BafA1-sensitive) to degrade the capsid ^25,26^. Treatment of NS2B/3-expressing cells with BafA1 to inhibit lysosomal degradation increased expression of NS2B/3 when expressed alone but did not rescue the relative loss of NS2B/3 in the presence of rhTRIM5 (Fig. 4h,i). This was despite the BafA1-sensitive rescue of p62/SQSTM1 which is a reported co-factor to TRIM5-mediated retrovirus restriction ^27^ (Fig. 4h). Selective autophagy of the HIV-1 capsid by TRIM5α is also mediated by Beclin, ATG5, p62, GABARAP and LC3 ^26^, but siRNA-mediated knockdown of these genes did not significantly relieve LGTV restriction (Supplementary Fig. 5a,b,c). Finally, the C-type lectin langerin, but not DC-SIGN, was previously shown to be sufficient for autophagic degradation of HIV-1 capsid by hTRIM5α ^25^. However, while DC-SIGN augmented LGTV replication as expected in its role as a flavivirus attachment factor ^28^, langerin expression had no effect and did not further increase the restriction of LGTV in TRIM5α expressing cells (Supplementary Fig. 5d), strongly suggesting that selective autophagy following virus entry or establishment of viral replication complexes is not the main mechanism of restriction. In contrast, treatment with epoxomicin (Fig. 4h,i) recovered the majority of NS3 in the presence of rhTRIM5α implicating proteasomal degradation of NS2B/3. This was supported by reciprocal IP of NS2B/3 ectopically co-expressed with rhTRIM5 in the presence of epoxomicin demonstrating a) increased interactions between TRIM5α and both the uncleaved NS2B/3 precursor and the mature, autocleaved NS3 protein, and b) increased ubiquitination of NS2B/3 co-precipitating with TRIM5α (Fig. 4g). TRIM5α did not appear to affect protease activity as autocleavage to produce NS3 measured by the ratio of NS2B/3:NS3 did not change in the presence of TRIM5α (Supplementary Fig. 5e). Furthermore, overexpression of K48R-HA ubiquitin (Ub) that cannot make K48-linked Ub chains, but not K63R-HA Ub, rescued expression of both NS2B/3 and rhTRIM5α (Fig. 4j), further suggesting that NS2B/3 degradation involves K48-linked ubiquitination which generally involves the proteasome.

**Fig. 5.**
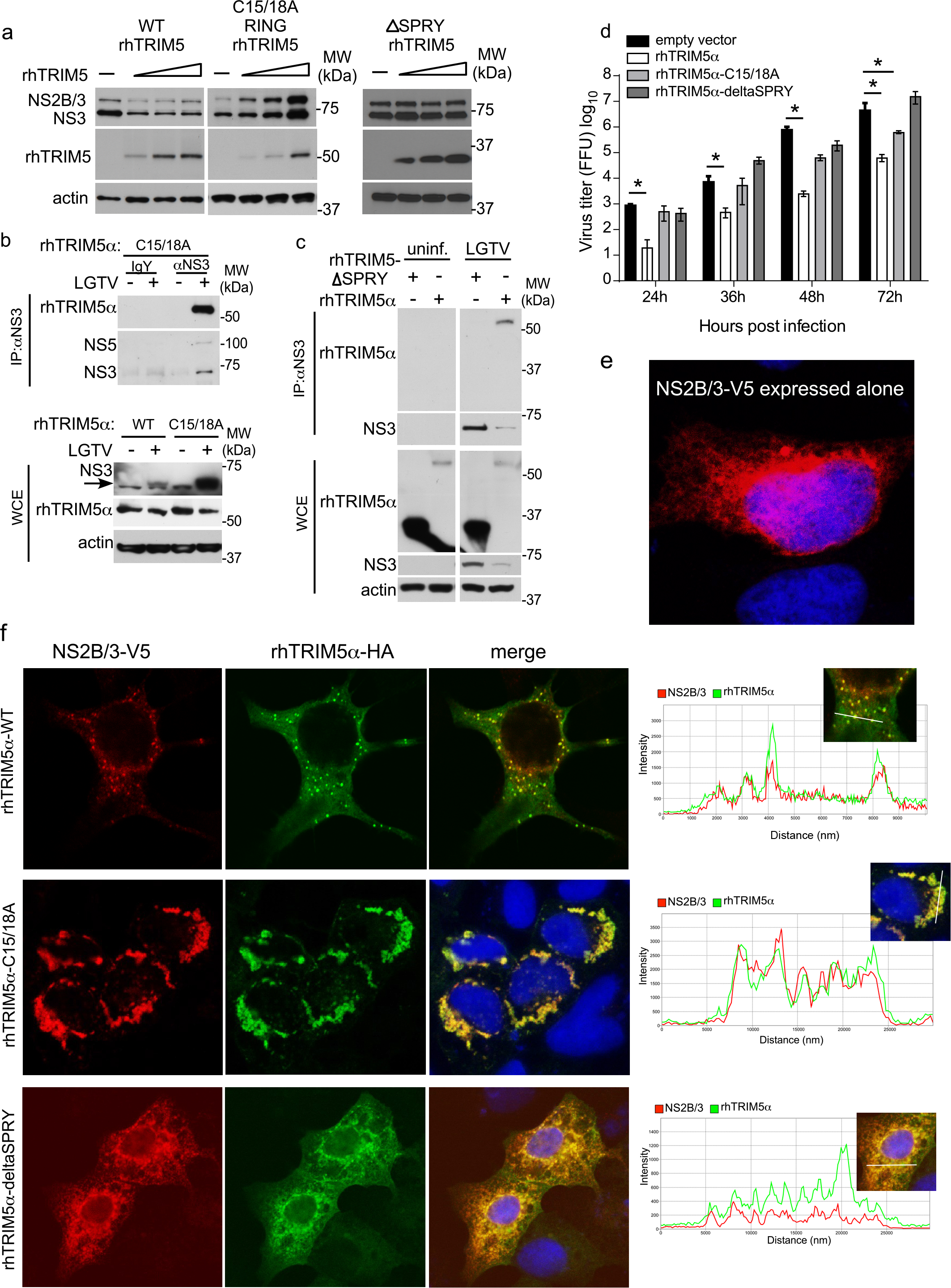
TRIM5α interaction with the flavivirus protease is associated with virus restriction. **a**, Western blot analysis following transfection of constant amounts of LGTV NS2B/3-V5 plasmid with increasing amounts of rhTRIM5α-HA, RING mutant rhTRIM5(C15/C18A)-HA, or rhTRIM5-delta SPRY-HA as indicated in HEK293 cells. **b-c**, Immunoprecipitation of NS3 from LGTV-infected HEK293 cells (MOI 0.01; 48 hpi) stably expressing **b**, RING rhTRIM5(C15/C18A)-HA or **c**, rhTRIM5α-HA or rhTRIM5-delta SPRY-HA. **d**, LGTV replication kinetics in HEK293 cells stably expressing rhTRIM5α-HA, RING mutant rhTRIM5(C15/C18A)-HA, rhTRIM5-delta SPRY or the empty vector control following infection at MOI of 0.001. All data are from three independent experiments (mean ± s.d., *P < 0.05 Mann Whitney test). **e-f**, HEK293 cells were co-transfected with LGTV NS2B/3-V5 (shown expressed alone in **e**), WT rhTRIM5α-HA, RING mutant rhTRIM5(C15/C18A)-HA or rhTRIM5-delta SPRY-HA. Slides were fixed and processed for indirect immunofluorescence staining with antibodies specific for HA (green) and V5 (red), and nuclei were counterstained with DAPI (blue). Images were analyzed using confocal microscopy with fluorescence intensity profiles measured across the white line of insets to demonstrate colocalization using Zen Imaging software.

To determine the domain of NS2B/3 recognized by TRIM5α, degradation assays were performed on various truncated NS2B/3 constructs (Fig. 4k). LGTV NS3 expressed without NS2B (Supplementary Fig. 5f) or the NS3 helicase domain alone (Supplementary Fig. 5g) was not sufficient for TRIM5α-mediated degradation. A construct containing the entire NS2B protein fused to the NS3 protease domain (NS2B-NS3pro) was also not degraded suggesting that NS2B alone is not sufficient as a target (Supplementary Fig. 5h). However, expression of NS3pro containing the 40 amino acids of NS2B required for NS3 protease activity in frame with a flexible glycine linker, the NS3 protease domain and the linker sequence between the NS3 protease and helicase domains, enabled degradation (Fig. 4l). Thus, the target for TRIM5α degradation requires NS2B in addition to NS3 sequences (NS3pro). Recognition of NS2B/3 is therefore likely dependent on protease conformation, but is independent of protease activity as the S138A active site mutant of NS2B/3 was also degraded (Supplementary Fig. 5i).

### TRIM5α interaction with the flavivirus protease is associated with virus restriction

The N-terminus of TRIM proteins is composed of a RBCC motif which includes a really interesting new gene (RING) domain, one or more B-box domains and a coiled-coiled (CC) domain ^29^. The RING and B-box can mediate conjugation of Ub thereby functioning as an E3 Ub ligase, whereas the CC domain allows oligomerization of TRIM proteins and formation of cytoplasmic bodies ^30^. The specificity of TRIM proteins is mainly determined by their C-terminal B30.2/SPRY domain that is responsible for binding to specific substrates including retroviral capsids ^15^. The C15/18A RING mutant of rhTRIM5α did not degrade NS2B/3 (Fig. 5a) and instead stabilized it consistent with retention of binding (Fig. 5b). Restriction of infectious virus production was also dependent on rhTRIM5α RING function, particularly at early times post infection (Fig. 5d). Compared to co-expression of LGTV NS2B/3 with WT-rhTRIM5α-HA, the C15/18A RING mutant retained strong colocalization by IFA, but lost the ability to form discrete cytoplasmic bodies (Fig. 5e,f). In contrast, deletion of the B30.2/SPRY domain eliminated degradation of NS2B/3 (Fig. 5a) associated with failure to bind NS3 in infected cells (Fig. 5c), reduced colocalization with ectopically expressed NS2B/3 (Fig. 5f), and the loss of antiviral activity (Fig. 5d). Importantly, this data directly links TRIM5α binding and degradation of NS2B/3 to its antiviral restriction capacity in the context of flaviviruses.

In the context of retroviruses, capsid binding by cyclophilin A (CypA) is required for virus replication ^31,32^ and substitution of the B30.2/SPRY domain of hTRIM5α with CypA facilitates hTRIM5α binding to HIV-1 and virus restriction ^29,33^. The tick-borne flaviviruses, including LGTV, are sensitive to Cyp inhibition (Supplementary Fig. 6a) ^34^, and CypA specifically is required for efficient virus replication (Supplementary Fig. 6b). However, while substitution of Owl Monkey CypA ^35^ or human CypA ^32^ for the hTRIM5α B30.2/SPRY domain suppressed replication of VSV-G pseudotyped HIV-1 (Supplementary Fig. 1a,b), these fusion proteins had no effect on replication of LGTV (Supplementary Fig. 6c, d). Thus, although CypA is required for flavivirus replication, and binds to nonstructural proteins NS5 ^36^ and NS4B ^37^ within viral replication complexes, TRIM5-CypA fusion proteins are not sufficient to restrict tick-borne flavivirus replication, confirming the importance of the B30.2/SPRY domain of TRIM5α in flavivirus restriction.

**Fig. 6:**
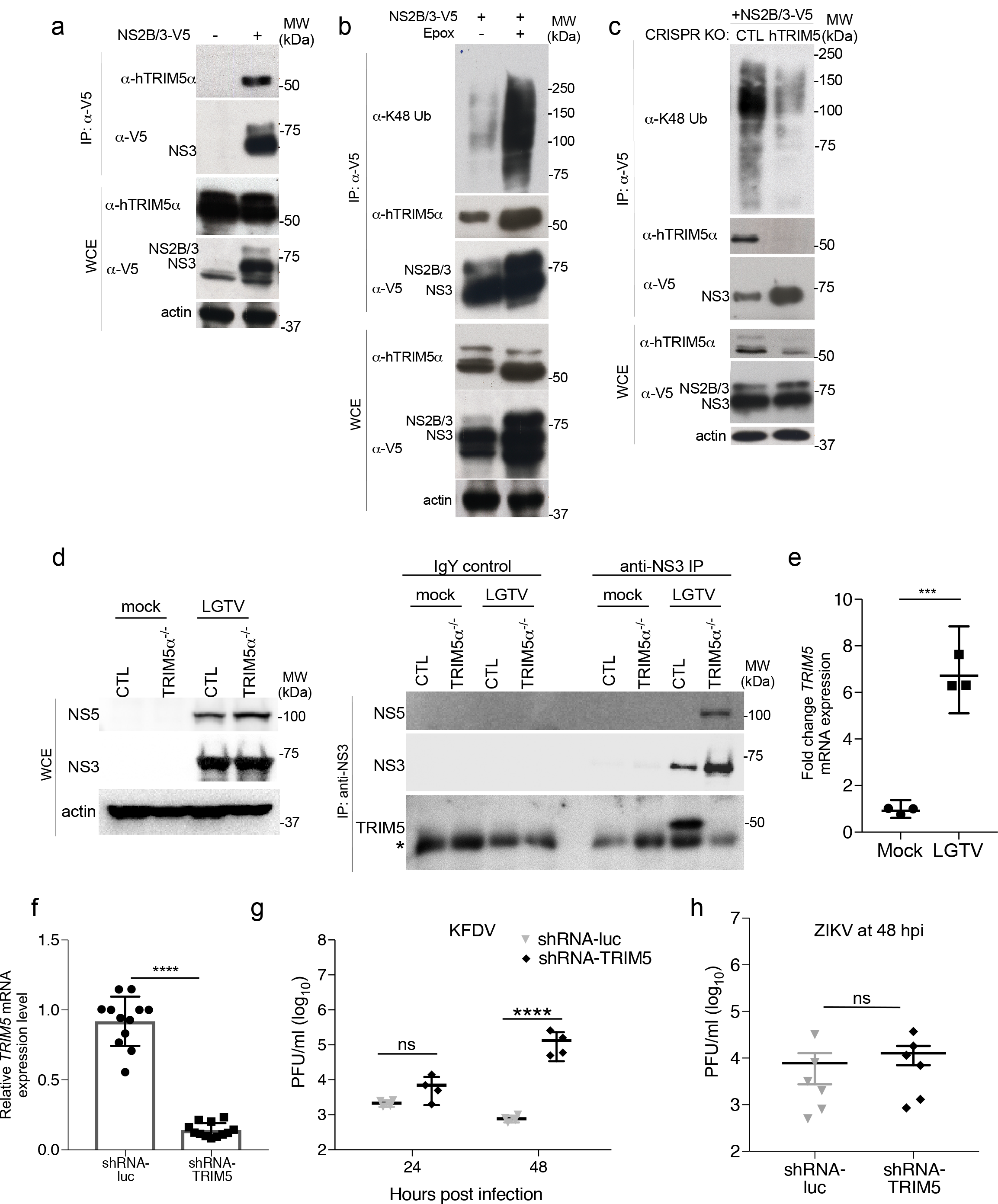
Endogenous human TRIM5 is an antiviral restriction factor for flaviviruses. **a**, IP of LGTV NS2B/3-V5 following ectopic expression in unmodified HEK293 cells and probed for TRIM5. **b**, IP of LGTV NS2B/3-V5 following ectopic expression and epoxomicin treatment in unmodified HEK293 cells. Western blots were probed for TRIM5 and K48-linked ubiquitin. **c**, IP of LGTV NS2B/3-V5 following ectopic expression in HEK293 cells transfected with plasmids encoding TRIM5 gRNA and Cas9. Western blots were probed for TRIM5 and K48-linked ubiquitin. **d**, HAP1 control and TRIM5-/-cells were infected with LGTV (MOI 0.1) and NS3 was immunoprecipitated at 48 hpi. Western blots were probed for TRIM5, NS3 and NS5. **e,f**, TRIM5 mRNA expression in primary human MDDCs **e**, infected with LGTV (MOI 5 at 24 hpi) or **f**, transduced with lentiviruses expressing shRNA-Luc (control) or shRNA-TRIM5 (mean ± s.d. from 3 experiments, *P < 0.0001 T-test). **g**, KFDV titers or **h**, ZIKV titers following infection of human MDDCs generated in part F. (MOI 0.1; mean ± s.d. from one of two experiments performed, ****P < 0.0001 ANOVA with Sidak post-test).

### Endogenous human TRIM5 is an antiviral restriction factor for flaviviruses

The role of human TRIM5α in suppression of HIV-1 has been controversial, in part because early studies suggested no restriction of laboratory strains of HIV-1. However, recent studies suggest that cytotoxic T lymphocyte (CTL)-selected HIV-1 isolates from so-called ‘elite controllers’ are susceptible to restriction by human TRIM5α ^38^, and genetic studies suggest that human polymorphisms in *TRIM5* impact disease progression ^13^. To further examine whether TRIM5α in human cells restricts flavivirus replication, we first immunoprecipitated LGTV NS2B/3 following ectopic expression in unmodified HEK293 cells which revealed an interaction with endogenous TRIM5α (Fig. 6a). Treatment of these cells with epoxomicin increased the levels of co-precipitating TRIM5 and NS2B/3 as well as the presence of endogenous K48-linked Ub smears in the complex (Fig. 6b), whereas depletion of TRIM5α by CRISPR/Cas9-mediated gene editing both increased levels of NS3 and decreased endogenous K48-linked Ub smears in the precipitates (Fig. 6c). Endogenous interactions between NS3 and TRIM5α were also confirmed in the HAP1 cells knocked out for TRIM5α by CRISPR/Cas9 and infected with LGTV (Fig. 6d). Finally, infection of primary human monocyte derived dendritic cells (DCs) resulted in upregulation of *TRIM5* expression (Fig. 6e). Silencing of *TRIM5* expression in human DCs by lentivirus-delivered shRNA expression ^39^ increased release of infectious KFDV by approximately 170 fold at 48 hpi compared to cells expressing shRNA specific for luciferase as a control (Fig. 6f,g). No effect of TRIM5α silencing was observed following infection with ZIKV (Fig. 6h). Together, these data demonstrate that human TRIM5α is a bona fide restriction factor for specific flaviviruses that functions through interactions with the viral replication complex and proteasomal degradation of NS3.

## Discussion

TRIM5α functions as an intrinsic cellular restriction factor that recognizes retrovirus capsids with high specificity and with definitive consequences for primate susceptibility to HIV-1 infection ^9–13,40,41^. Our work significantly extends the paradigm of TRIM5α as an antiviral restriction factor and suggests that, in contrast to the current view, TRIM5α exhibits a remarkable plasticity in recognition of unrelated viruses. Both human and rhesus TRIM5α are capable of restriction of specific flaviviruses within the TBEV serocomplex, and endogenous TRIM5α is required for the antiviral effects of type I IFN against sensitive flaviviruses in human cells. Interestingly, a recent report revealed that IFN-dependent activation of the immunoproteasome in human CD4^+^ T cells enables K48-ubiquitin-dependent, TRIM5α-mediated, restriction of HIV-1^42^. Our results suggest that IFN is not required for TRIM5α to degrade NS2B/3 via the proteasome, although restriction of LGTV and TBEV by TRIM5α in human cells was strongly evident when cells were pre-treated with IFN. Thus, it will be important to determine the interplay between IFN, TRIM5α, and flavivirus restriction. The mechanisms by which some flaviviruses evade restriction is unknown, and could involve evolution to avoid TRIM5α recognition at the sequence level, or more direct antagonism of TRIM5α and its putative cellular partners that may regulate this process. Alternatively, given the role of IFN for TRIM5α-mediated restriction of HIV-1 in human cells, the varied IFN antagonism strategies utilized by flaviviruses^6^ may aid in TRIM5α escape.

The rapid evolution of the TRIM5 gene throughout primate evolution is associated with selection pressure from lentivirus capsid sequences ^16^. It is therefore unclear how evolutionary selection of TRIM5α for retrovirus restriction has left the protein with enough flexibility to maintain antiviral activity against flaviviruses. It may be possible that ancient flavivirus-like viruses have influenced the evolution of human TRIM5α. However, the timeframe of flavivirus evolution is in the order of thousands of years in contrast to millions of years for retroviruses and the TRIM5 gene ^14^. The *Flaviviridae* includes the more ancient genera of Hepaciviruses, although evidence for a zoonotic origin of hepatitis C virus (HCV) in non-human primates is not strong despite the extremely narrow host range of HCV limited to humans and chimpanzees ^43^. Therefore, it seems unlikely that flaviviruses influenced positive selection of the *TRIM5* gene within the human lineage. However, our work raises the possibility that human polymorphisms within the *TRIM5* locus could influence resistance to infection with medically important flaviviruses. Thus, understanding the genetic trade-offs in both TRIM5α and NS2B/3 that enable restriction of flaviviruses versus retroviruses represents an important model to illustrate how host resistance is shaped by multiple pathogens, and might provide new insight to human susceptibility to emerging flaviviruses. The finding that primate TRIM5α can recognize and degrade NS2B/3 from specific flaviviruses combined with a strong antiviral role in the type I IFN response suggests that TRIM5α has high potential to function as an important human barrier to infection with emerging flaviviruses. We speculate that viral resistance to TRIM5α-mediated restriction may be an important factor in enabling use of primates, including humans, as reservoirs for viruses like YFV, DENV and ZIKV.

## Acknowledgements

This work was supported by the Division of Intramural Research of the National Institute of Allergy and Infectious Diseases (NIAID), National Institutes of Health (NIH), by NIH grants R01AI111809 and DP1DA034990 to J.L., and RO1AI137011 and DP1DA046108 to S.L.S. S.L.S. is a Burroughs Wellcome Investigator in the Pathogenesis of Infectious Disease. N.R.M. was supported by a graduate fellowship from the NSF, and a PDEP award from the Burroughs Wellcome Fund.

## Contributions

Conceptualization, A.I.C and S.M.B.; Methodology, A.I.C., N.R.M, S.L.S., and S.M.B.; Investigation, A.I.C., N.R.M., K.L.M., R.M.B., V.R.M., O.M.-S., S.J.R., F.B., G.L.S., K.J.L., V.N., B.H.Y., R.T.T., K.K., and S.M.B.; Resources, R.T.T., S.L.S. and S.M.B.; Data Curation, A.I.C., N.R.M., S.M.B.; Writing – Original Draft, A.I.C. and S.M.B.; Writing – Review & Editing, A.I.C., N.R.M., R.T.T., S.L.S. and S.M.B.; Resources, J.L., V.M.H, R.T.T., S.L.S. and S.M.B.; Visualization, A.I.C., N.R.M., V.R.M., K.L.M., O.M.-S., S.J.R., G.L.S., K.J.L., V.N., S.L.S. and S.M.B.; Supervision, J.L., R.T.T., S.L.S. and S.M.B.

## Methods

### Cell Culture and Reagents

HEK293T cells (human embryonic kidney, ATCC; CRL-3216), HEK293 cells (human embryonic kidney, ATCC; CRL-1573), CRFK cells (feline kidney, ATCC; CCL-94), A549 cells (lung carcinoma, ATCC; CCL-185) and Vero cells were cultured in Dulbecco’s modified Eagle media (Gibco; 11995) supplemented with 10% fetal bovine serum (Gibco; 16000-044), 2 mM L-glutamine (Invitrogen; 25030-081), and 1% antibiotics (Gibco; 15140) (complete media) at 37°C and 5% CO_2_. Cell culture grade epoxomicin and MG132 (proteasomal inhibitors), bafilomycin A1 (Baf-A1), puromycin and blasticidin were purchased from Sigma. Interferon β (IFNβ) was purchased from PBL Assay Science (#11410-2).

### Virus Infections and Lentivirus production

The viruses used in this study were handled under biosafety level 2 (BSL2), BSL3 and BSL4 conditions at the Rocky Mountain Laboratories Integrated Research Facility in accordance with DSAT regulations for study of select agents and Institutional Biosafety approvals (Hamilton, MT). The viruses in this study include: Langat virus (LGTV) strain TP21 (from Dr. A. Pletnev, NIAID, NIH), TBEV strain Sofjin (also referred to as Russian spring summer encephalitis [RSSE] virus), Kyasanur forest disease virus (KFDV) [from Dr. M. Holbrook, NIAID, NIH], Powassan virus (POWV, strain LB) and West Nile virus (strain NY99) [from the WRCEVA], Dengue virus (DENV-2, strain New Guinea C) from Dr. Adolfo García-Sastre), Zika virus (ZIKV, strain 2013 French Polynesia, from Dr. David Safronetz) and Yellow fever virus (YFV, strain 17D), from NIH Biodefense and Emerging Infections Research Resources Repository, NIAID, NIH, NR115. All viruses were propagated as previously described ^44^. Cell monolayers were infected for 1 h at 37°C, after which virus inoculum was removed and cells replenished with fresh cell culture medium. Virus titers are represented as plaque forming units (PFUs) or focus forming units (FFU) per 1 ml.

HIV-1 virus pseudotyped with VSV-G and encoding a GFP reporter for single-cycle infection assays were packaged in 293T cells seeded at a concentration of 1×10^6^ cells/well in a 6-well dish. One day after seeding, cells were co-transfected with 0.5 μg pMDLg/pRRE, 0.25μg pRSV-Rev, 0.2μg pMD2.G, and 1μg pRRLSIN.cPPT.PGK-GFP.WPRE (plasmids 60488, 12253, 12252 respectively available from Addgene). Cells were transfected using TransIT-293 at a 1:3 ratio (μg DNA:μl TransIT-293). After 48 hours, supernatant containing viruses was harvested, filtered, and frozen. For infection assays, CrFK stable cells lines were plated at a concentration of 7.5×10^4^ cells/well in a 24-well plate or HEK293 stable cell lines were plated at a concentration of 1.0×10^5^ cells/well in a 24-well plate, and infected with HIV-1 single-cycle virus. Two days post-infection, cells were fixed, washed, resuspended in PBS supplemented with 1% FBS, and analyzed by flow cytometry for expression of GFP using the BD Bioscience Fortessa cell analyzer.

### Lentivirus generation expressing shRNAs

The shTRIM5 and shluciferase lentiviruses were generated by transfecting HEK293T cells with lentivirus shRNA plasmid (pAPM CoE D4 L1221 or pAPM CoE D4 TRIM5 ts2 for shluciferase or shTRIM5, respectively), pSPAX2, and pMD.G using the ProFection Mammalian Transfection System (Promega). pAPM CoE D4 is a truncated derivative of the pAPM lentiviral vector that expresses the puromycin acetyltransferase and miR30-based shRNA from the SFFV promoter (Pertel et al. 2011). The target sequences are: pAPM CoE D4 L1221 5ʹ-TACAAACGCTCTCATCGACAAG-3ʹ and pAPM CoE D4 TRIM5 ts2 5ʹ-TGCCAAGCATGCCTCACTGCAA-3ʹ. The vpx-vlp was generated by transfecting 293T cells with pMD.G and SIV_MAC_ packaging plasmid kindly provided by Dr. Andrea Cimarelli ^45^. Media was replaced 18-20 hours post transfection (hpt). Supernatant was harvested at 48 hpt, passed through a 0.45 um filter, and ultracentrifuged over a cushion consisting of 25% sucrose in TNE buffer (10 mM Tris-HCl, pH 7.5, 1 mM EDTA, 100 mM NaCl, pH 7.4) at 28,000 rpm in a SW-28 Rotor (Beckman). Lentivirus pellets were resuspended in PBS, aliquoted, and stored at −80°C prior to use. shRNA-luc and shRNA-TRIM5 lentivirus titers were normalized by serial dilution on HEK293 cells followed by puromycin selection.

### Knockdown of TRIM5 in Human monocyte-derived dendritic cells (hMDDC) cultures

Human monocyte cultures ^46^ were seeded in 48-well plates and transduced with a combination of vpx-vlp and shControl or shTRIM5 lentivirus for three hours followed by addition of IL-4 and GM-CSF-conditioned RPMI media. Conditioned media was replenished at 3 days post transduction (dpt). Five dpt, cells were collected to confirm knockdown of TRIM5 transcripts by qRT-PCR. Remaining cells were infected with ZIKV PRABC59 (MOI = 5) or KFDV (MOI = 0.1) for 48 hours. Supernatants were collected at the indicated times, and virus was measured in the supernatant by limiting dilution plaque assay.

### Expression constructs

HA-tagged (C-term) human and rhesus *TRIM5* in the pLPCX retroviral vector were obtained from the National Institutes of Health AIDS Research and Reference Reagent Program. HA-tagged (C-term) owl monkey *TRIM-CypA* in the pLPCX retroviral vector was a kind gift from Dr. Michael Emerman (Fred Hutchinson Cancer Research Center). Approximately 5×10^6^ HEK293 cells were used to isolate RNA with the All Prep RNA/DNA Mini Kit (Qiagen; 80204). cDNA was generated using 1μg of RNA with oligo(dT) primers and the Superscript III First-Strand Synthesis System (Invitrogen; 18080-051). This cDNA was used as a template to amplify the *CypA* coding region (see below). All primers used in this study for qRT-PCR or to generate constructs, along with a description of their use, can be found in Extended Data Table 1. Human *TRIM22* was amplified from a pcDNA3 construct kindly provided by Dianne Lou. *TRIM-CypA and TRIM-RanCyp* constructs were generated by amplifying fragments (aa 1-309 from human *TRIM5* in pLPCX and the complete coding sequence of *CypA* from HEK293 cDNA) with 20-25bp overlapping regions. Overlapping fragments were spliced together in a PCR reaction using each fragment as a template and outside flanking primers. Human and rhesus *TRIM5delB30.2* constructs were generated using pLPCX templates and primers that amplify aa 1-276 from human *TRIM5* or 1-278 from rhesus *TRIM5.* All above PCR reactions were carried out using PCR Supermix High Fidelity (Thermo Fisher; 10790020) with an annealing temperature of 58°C. Constructs were TA-cloned into the gateway entry plasmid pCR8 (Invitrogen; K2500-20). An LR Clonase II reaction (Invitrogen; 11791-100) was used to move these constructs into a Gateway-converted pLPCX retroviral packaging vector (Clontech; 631511). The RING C15/18A mutant of *TRIM5* was generated using PfuTurbo DNA polymerase (Stratagene; 600250) with an annealing temperature of 55°C. Parental pLPCX plasmids were used as a template along with primers containing the mutations of interest. Constructs expressing LGTV and WNV_NY99_ NS2B/3 and NS5 were generated as previously described ^44^. Expression plasmids for Langerin (HG13040-UT) and DC-SIGN (HG10200-UT) were purchased from Sino Biological.

### Generation of stable cells lines

To make cell lines that stably express *TRIM5* constructs, pLPCX retroviral vectors were used to transduce HEK293 cells. To generate the retroviruses used for transduction, HEK293T cells were seeded at a concentration of 1×10^6^ cells/well in a 6-well dish. 24 hours later each well was transfected with 2 μg pLPCX construct (empty or encoding the gene fragment of interest), 1 μg pCS2-mGP encoding MLV gag-pol^2^, and 0.2 μg pC-VSV-G (provided by Hyeryun Choe) at a final 1:3 ratio of DNA to TransIT-293 (μg DNA: μl TransIT-293). Supernatants were collected after 48 h, passed through a 0.2 μm filter, and used to infect HEK293 cells grown in complete media. HEK293 cells were seeded in a 12-well dish at a concentration of 7.5×10^4^ cells/well. After 24 h, varying amounts of retrovirus from each construct were added to cells along with polybrene (Sigma; 107689) at a final concentration of 10 μg/mL. After 24 h, media containing 0.75 μg/ml puromycin (Sigma; P8833) was added to select for transduced cells. Cell lines were eventually expanded into 10 cm dishes, checked for expression of the appropriate construct by Western blot, and frozen down in 1 mL aliquots containing complete media supplemented with an additional 10% FBS (total of 20%) and 5% DMSO. A549 cells were stably knocked-down using lentiviruses coding short hairpin RNAs (shRNAs) against *Cyclophilin A*, *B* and *non-targeting* (control) *as* previously described (kindly provided by Prof. Ralf Bartenschlager)^47^. HAP1 cells edited within the TRIM5 gene were generated by Horizon Genomics (Vienna) with the RNA guide sequence: CGATTAGGCCGTATGTTCTC.

### Antibodies

HA-tagged constructs for western blotting were detected using a 1:5000 dilution of anti-HA-peroxidase antibody (Roche clone 3F10, #12013819001). HA-tagged constructs for indirect immunofluorescence were detected using anti-HA (Zymed, #71-5500). β-actin was also detected as a loading control using a 1:10,000 dilution of mouse anti-β-actin (Sigma, A5441). A 1:3,000 dilution of goat anti-mouse (Dako, #P0447), anti-rabbit (Thermo Scientific, #P0448) or anti-chicken (Millipore, #12-341) horseradish peroxidase-conjugated antibody was used as a secondary probe. V5 tagged constructs were probed with anti-mouse V5 (Invitrogen #R960-25). Blots were developed using the ECL Plus detection reagent (GE Healthcare, #RPN2132). Antibodies to detect viral antigens, LGTV (NS3 and NS5) (previously described in Taylor et al., 2011), WNV-NS3 (R&D Systems, #AF2907) and dsRNA antibody J2 (English& Scientific Consulting, #10010200). Autophagy and cellular markers were detected using LC3B (Nanotools, #5F10), GABARAP (Cell Signaling, #E1J4E), Beclin-1 (Novus Biologicals, # 110-53818), ATG5 (Cell Signaling, #2630), p62 (BD Transduction Laboratories, #610833), cyclophilin A (Enzo, #BML-SA296-0100), cyclophilin B (Thermo Scientific, #PA1-027A), langerin (R&D Systems, #AF2088) and DC-SIGN (BD Biosciences, #551186).

### Immunoprecipitation (IP) and Western Blot Analysis

293 cells were washed three times with PBS (1X) and lysed on ice in RIPA buffer (50 mM Tris-HCl [pH 7.6], 150 mM NaCl, 0.1% SDS, 1% Igepal, and 0.5% Na-deoxycholate) with protease inhibitor cocktail (Roche). For IPs of over-expressed proteins, 2 wells of a 6 well dish at 1×10^6^ cells/well were used per reaction; for IPs of virus-infected stable TRIM5 HEK293 cells, a 10cm dish of 7×10^6^ cells/dish was used per reaction; for detection of endogenous TRIM5, HEK293 or HAP1 cells were grown to confluency in 3-4 T150 tissue culture flasks. Samples were subjected to centrifugation for 10 min at maximum speed to remove cellular debris. Protein G-conjugated agarose beads (Roche) or PrecipHen for chicken antibodies (Aves Labs) were used to clear cell lysates at 4°C for 3 h. Samples were centrifuged to remove beads, and 2 μg of antibody analogous to the protein of interest was added to each lysate for 1 h with rotation at 4°C. 50 μl protein G-agarose or PrecipHen beads and were incubated with rotation at 4°C overnight. Lysates were subjected to centrifugation, and beads were washed three times with RIPA buffer prior to elution by incubation at 95°C in 1× sample buffer (62.5 mM TRIS [pH 6.8], 10% glycerol, 15 mM EDTA, 4% 2-ME, 2% SDS, and bromophenol blue). For western blot analysis HEK293 cell lines were grown to confluency in a 12-well or 6-well dish, collected using a cell scraper, and lysed in RIPA buffer containing complete protease inhibitor (Roche, #11836170001). After quantification of protein concentration using a Bradford assay, 30 μg of whole cell extract was resolved using a 10% polyacrylamide gel and transferred to a nitrocellulose membrane. Ubiquitination assays were performed as previously described (Campbell et al., 2015). Densitometry analysis was performed using ImageJ software.

### Confocal Microscopy

Cells were seeded onto 4 well Lab-Tek II chamber slides overnight. Slides were prepared by washing cells twice with PBS (1X) and subsequently fixed with paraformaldehyde (4%) for 10 min. For double-stranded RNA (dsRNA) staining, cells were fixed with methanol (100%) for 5 min at −20°C. Slides fixed with paraformaldehyde (4%) were further incubated with permeabilization buffer (Triton X-100 [0.1%], sodium citrate [0.1%]) for 5 min at room temperature and incubated with blocking buffer (PBS[1X], BSA [0.5%] and goat serum [1%]) for 30 min. Cells were incubated with primary antibody overnight at 4°C, washed three times with PBS (1X) and further incubated with secondary antibody conjugated to Alexa-488, −594 or −647 (Molecular Probes) for 1 h. Slides were washed three time with PBS(1X) and once with ddH_2_0, and mounted onto glass coverslips using Prolong Gold + DAP1 mounting media (Molecular Probes). Processed slides were imaged using a Zeiss LSM710 confocal microscope and vector profiles analyzed using Zen software (Carl Zeiss).

### Flow cytometry

Cells were harvested at 48 hpi and processed for flow cytometry analysis. Cells were stained with LIVE/DEAD Fixable Aqua Dead Cell Stain Kit (ThermoFisher) and fixed with 4 % paraformaldehyde for 20 min at RT. Cells were permeabilized with saponin-containing buffer and probed with anti-E 11H12 antibody. Data were generated using an LSRII flow cytometer (BD Biosciences) and analyzed using FlowJo (Tree Star).

### RNA Isolation and quantitative RT-PCR

RNA was isolated from cells using RNeasy kit (Qiagen) and genomic DNA was removed with RNase-free DNase (Qiagen). Reverse transcription of RNA was performed using Superscript Vilo cDNA Synthesis Kit (Invitrogen) according to manufacturer’s protocol. Details of TaqMan probes specific for *TRIM5*, hypoxanthine-guanine phosphoribosyltransferase (*HPRT*), interferon beta (*IFNβ*), interlukin −6 (*IL6*), tumor necrosis factor alpha (*TNFα*) and C-X-C motif chemokine 10 (*CXCL10*) are listed Extended Data Table 1. All probes were obtained from Applied Biosystems. Reactions for Real-time RT-PCR were set up in triplicate, cycled and data was collected on the Applied Biosystems GeneAmp 9500 Sequence detection system. Quantification of relative gene expression was relative to untreated controls with comparative C_T_ method.

### RNA interference

HEK293 and A549 cells were transfected with 15 pmol of siRNA using Lipofectamine RNAiMAX (Life Technologies). siRNAs (Dharmacon; SMART pool) were specific against TRIM5 (L-007100), LC3B (L-012846), GABARAP (L-012368), Beclin-1 (L-010552), ATG5 (L-004374) and p62 (L-010230).

### Statistical Analysis

All data were evaluated for significance using one-tailed unpaired Student’s *t-*test, or Mann-Whitney U test or one-way ANOVA with Tukey post-test using GraphPad Prism 7 software.

