## Supplemental Figures for "TRIM5α restricts flavivirus replication by targeting the viral protease for proteasomal degradation"

**Supplementary Figures**

**Supplementary Fig. 1. Human (h) and rhesus (rh) TRIM5α restrict flavivirus infection. a,** Western blot showing relative expression levels of primate TRIM5α-HA in stable cell lines used in this study including hTRIM5α, rhTRIM5α, hTRIM5-hCypA (human cyclophilin A), hTRIM5-OMCypA (Owl monkey cyclophilin A), hTRIM22 or empty vector (control). **b,** Cell lines in A. were infected with HIV-1 virus pseudotyped with VSV-G and encoding a GFP reporter and examine for infection at 48 hpi by flow cytometry for GFP. **c-d,** CrFK cells stably expressing hTRIM5α-HA or empty vector (control) were infected with **c.** TBEV or **d,** WNV at an MOI of 1 and infectious virus was assayed in supernatants by plaque assay at 24 h intervals. ND, none detected. Data shown from one of two experiments performed in triplicate (mean ± s.d.; $*P < 0.05$, $**P < 0.01$). **e,** Titers of LGTV at 48 hpi in empty vector- and rhTRIM5α-HA HEK293 cells following infection at MOI 10. **f,** Sequencing results of the *TRIM5* mRNA sequence from Hap1 cells transduced with CRISPR guide sequence for *TRIM5* and Cas9. The resulting cells had a deletion in nucleotides 182-215 of the mRNA resulting in a missense mutation. **g,** mRNA expression of *TRIM5*, *TRIM25* and *IFIT1* in Hap1 gene edited to knock-out TRIM5 expression ($TRIM5^{-/-}$) and treated with IFNβ for 6 h (mean ± s.d.; $*P < 0.001$).

**Supplementary Fig. 2. TRIM5α localizes to sites of viral RNA replication to mediate restriction. a,** Localization of NS3 (red) and dsRNA (green) in LGTV-infected cells. **b,** Localization of NS3 (red) and hTRIM5α (green) in mock- or LGTV-infected hTRIM5α-HA-expressing HEK293 cells. Nuclei were stained with DAPI (blue). Images were captured and analyzed using confocal microscopy. **c,** Stable rhTRIM5α-HA cells were infected with WNV (MOI 0.01) and stained for dsRNA (red) and rhTRIM5α (green) at 24 hpi. Nuclei were counterstained with DAPI. Inset shows the region indicated by a white box. **d,** Example of the intensity profile along vectors drawn through dsRNA staining in rhTRIM5α-HA cells infected with WNV. Asterisk indicates region of co-staining. **e,** rhTRIM5α-HA-expressing HEK293 cells were infected with WNV (MOI 0.01) and NS3 was precipitated at 48 hpi. No interacting rhTRIM5α-HA was detected.

**Supplementary Fig. 3. rhTRIM5α does not form cytoplasmic bodies at sites of dsRNA during replication of ZIKV or DENV. a,** Stable rhTRIM5α-HA cells were infected with LGTV (MOI 5), ZIKV (MOI 0.01) or DENV (MOI 0.01) and stained for dsRNA (red) and rhTRIM5α (green) at 24 hpi. Nuclei were counterstained with DAPI. Insets show the region indicated by a white box. **b,** Examples of the intensity profiles along vectors drawn through dsRNA staining in rhTRIM5α-HA cells infected with LGTV, ZIKV or DENV as well as quantification of rhTRIM5α-HA intensity changes in uninfected and infected cells. DENV and ZIKV have no obvious accumulation of cytoplasmic TRIM5 aggregates although TRIM5α signal is present at concentrations of dsRNA as measured by intensity across vectors and indicated by asterisks.

**Supplementary Fig. 4. TRIM5α does not strongly associate with LGTV NS5. a,** Reciprocal IP of LGTV NS5-V5 or rhTRIM5α-HA following co-expression in HEK293 cells. **b,** An example of the quantification of rhTRIM5α-HA aggregation measured by intensity across vectors drawn in stable rhTRIM5α-HA HEK393 cells that are untransfected (control) or in cells positive for LGTV NS2B/3-V5. The quantification for these images is summarized in Fig. 5b.

**Supplementary Fig. 5. TRIM5α does not serve as a selective autophagy receptor during flavivirus replication. a,** LGTV titers in control and rhTRIM5α-HA stable cell lines transfected with control non-targeting (NT) siRNA to demonstrate levels of restriction in these experiments without data normalization. **b,** Empty vector (control) and rhTRIM5α-HA stable expression cells were depleted for autophagy-related genes using specific siRNAs as indicated. At 48 h post silencing, cells were infected with LGTV (MOI 0.01) for another 48 h and virus titers from supernatants were determined using plaque assays. Data was normalized to percentage change in titer from cells treated with NT siRNA. All data are from 4-6 independent experiments performed in triplicate (ns, not significant). **c,** Western blot analysis for validation of siRNA knockdown 96 h post silencing using specific antibodies for BECLIN-1, ATG5, GABARAP, LC3B, p62 and b-actin in empty vector (EV) or rhTRIM5α-HA (T5) cell lines. **d,** rhTRIM5α-HA or hTRIM5α-HA stable cell lines were transfected with expression plasmids for GFP, DC-SIGN or Langerin. 24 h later, cells were infected with LGTV (MOI of 0.01) and virus titers were determined at 48 hpi by plaque assay. Data are shown from three independent experiments performed in triplicate (mean ± s.d., $*P < 0.05$, $**** P < 0.0001$, ns, not significant). **e,** Quantification by densitometry of the ratio of cleaved LGTV NS3: total NS2B/3 in the presence or absence of rhTRIM5α-HA across 10 independent experiments, suggesting no effect of TRIM5α on the protease activity of NS2B/3. **f-i,** Western blot analysis of HEK293 cells transfected with increasing amounts of rhTRIM5α-HA and constant amounts of **f,** LGTV NS3-V5, **g,** LGTV NS3 helicase domain, **h,** LGTV NS2B plus the NS3 protease domain, **i,** LGTV NS2B/3 S138A protease mutant. Lysates were probed specifically for HA, V5 and b-actin.

**Supplementary Fig. 6. TRIM5α restriction of flaviviruses is independent of its association with CypA. a,** A549 cells were infected with LGTV (MOI of 0.01) for 24 h followed by treatment with increasing amounts of alisporivir or DMSO (vehicle) for 48 h. Supernatants were harvested and virus titer determined using plaque assays. **b,** shRNA specific for cyclophilin A (CypA), cyclophilin B (CypB) or non-targeting (NT) control were transduced into A549 cells followed by LGTV infection (MOI of 0.01). Supernatants were harvested at indicated time points and virus titers were determined using plaque assay. Western blot inset shows specific knock down of CypA and CypB as compared to NT control. **c-d,** HEK293 cells stably overexpressing C-terminally HA tagged fusion constructs for **c,** hTRIM5-OMCypA-HA or **d,** hTRIM5-hCypA-HA were infected with LGTV (MOI of 0.01) and virus supernatants were harvested at indicated time points to determine virus titers by plaque assays (data is shown from one of two experiments performed in triplicate; mean ± s.d.) ($*P<0.05$; $**P<0.01$).

Figure S1

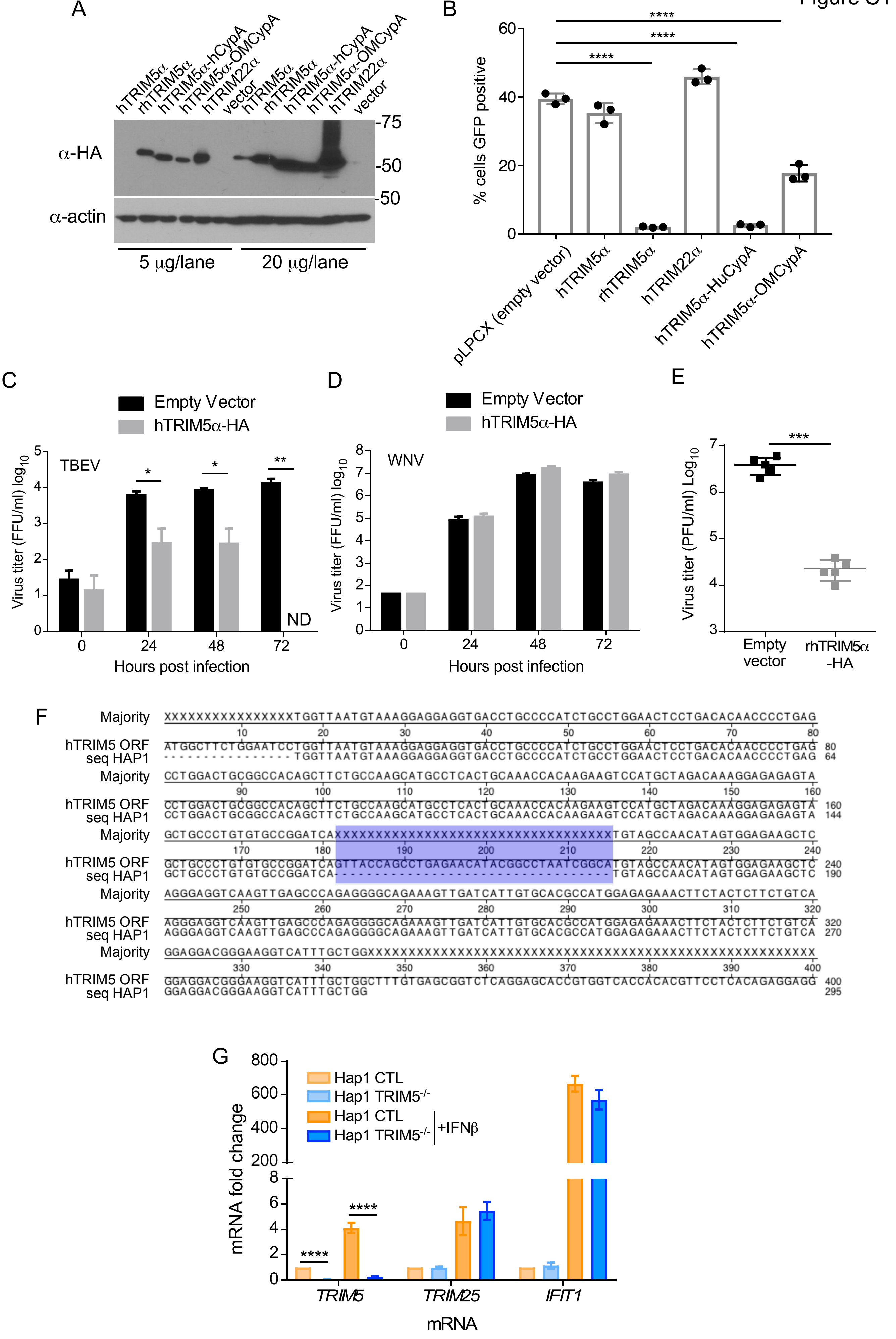

A
hTRIM5α
rhTRIM5α
hTRIM5α-hCypA
hTRIM5α-OMCypA
hTRIM22α
vector
-75
-50
-50
α-HA
α-actin
5 μg/lane
20 μg/lane
B
****
% cells GFP positive
pLPCX (empty vector)
hTRIM5α
rhTRIM5α
hTRIM22α
hTRIM5α-HuCypA
hTRIM5α-OMCypA
C
Empty Vector
hTRIM5α-HA
TBEV
ND
Virus titer (FFU/ml) log10
Hours post infection
D
WNV
E
Virus titer (PFU/ml) Log10
***
Empty vector
rhTRIM5α -HA
F
Majority
hTRIM5 ORF
seq HAP1
G
Hap1 CTL
Hap1 TRIM5-/-
+IFNβ
mRNA fold change
TRIM5
TRIM25
IFIT1
mRNA

Suppl Fig. 2

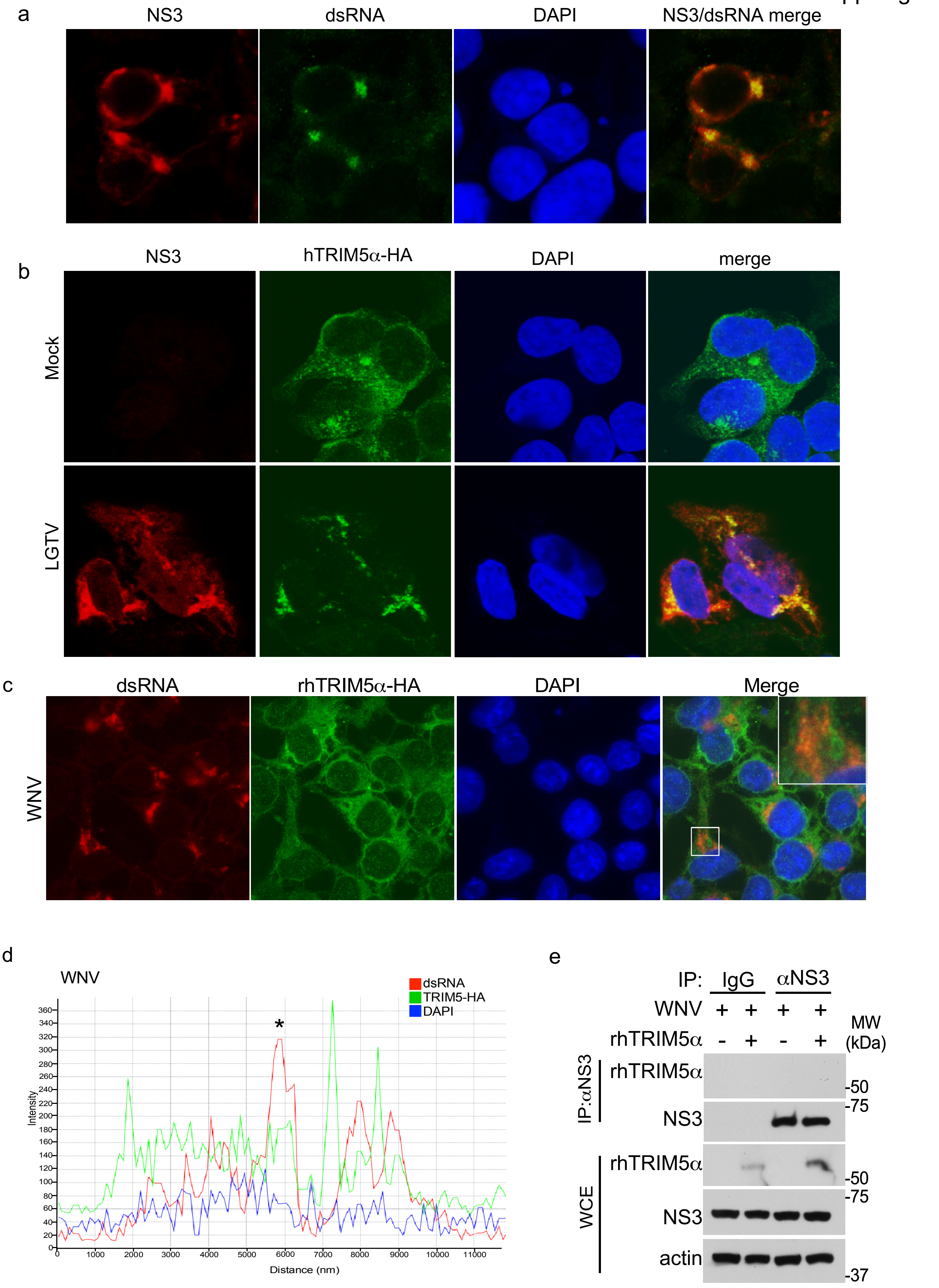
a
NS3
dsRNA
DAPI
NS3/dsRNA merge
b
NS3
hTRIM5α-HA
DAPI
merge
Mock
LGTV
c
dsRNA
rhTRIM5α-HA
DAPI
Merge
WNV
d
WNV
dsRNA
TRIM5-HA
DAPI
*
Intensity
Distance (nm)
e
IP: IgG αNS3
WNV + + + +
rhTRIM5α - + - +
MW (kDa)
IP:αNS3
rhTRIM5α
NS3
WCE
rhTRIM5α
NS3
actin
-50
-75
-50
-75
-37

Suppl. Fig. 3

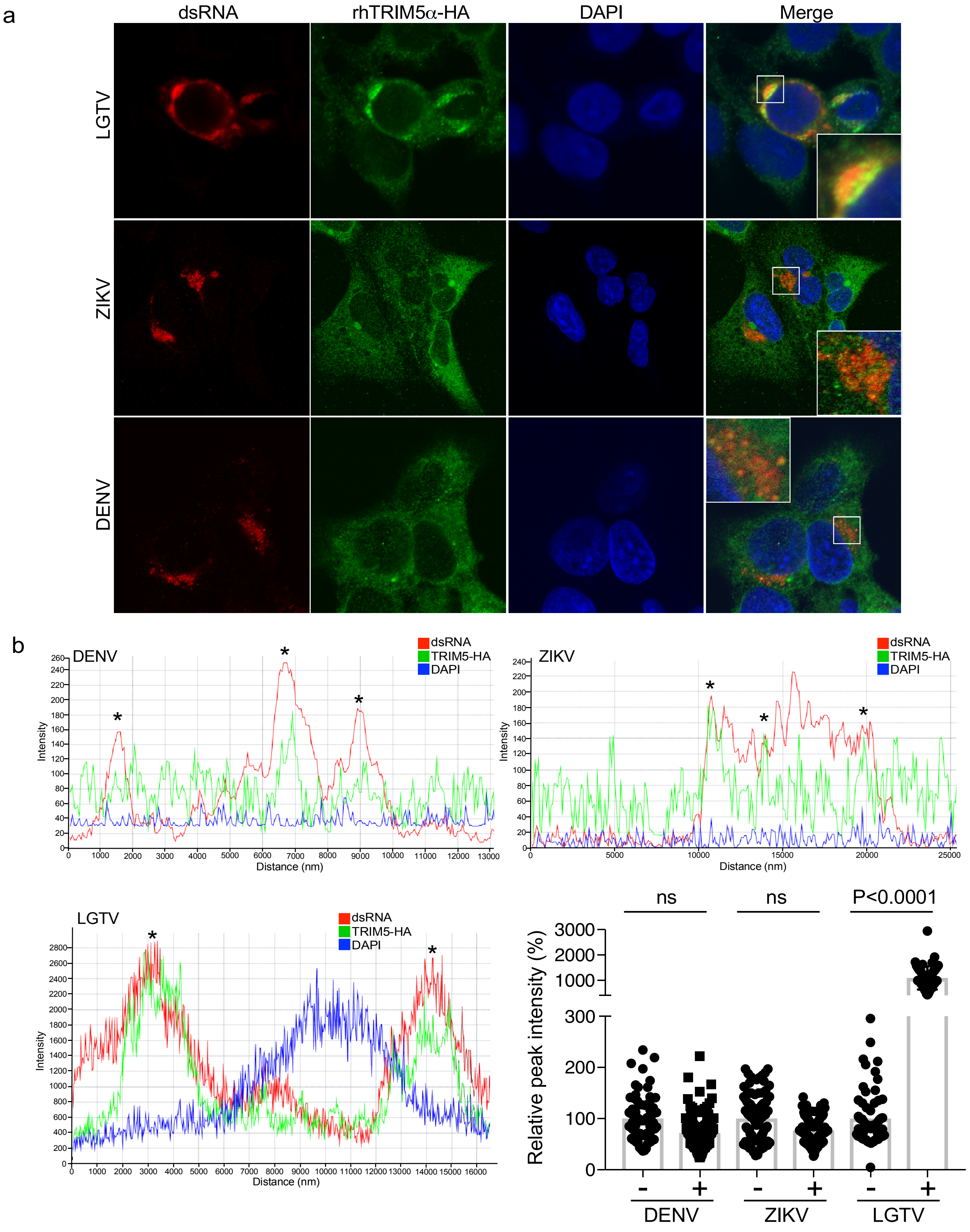

a
dsRNA
rhTRIM5α-HA
DAPI
Merge
LGTV
ZIKV
DENV
b
DENV
dsRNA
TRIM5-HA
DAPI
Intensity
Distance (nm)
*
ZIKV
dsRNA
TRIM5-HA
DAPI
Intensity
Distance (nm)
LGTV
dsRNA
TRIM5-HA
DAPI
Intensity
Distance (nm)
ns
ns
P<0.0001
Relative peak intensity (%)
3000
2000
1000
300
200
100
0
-
+
DENV
ZIKV
LGTV

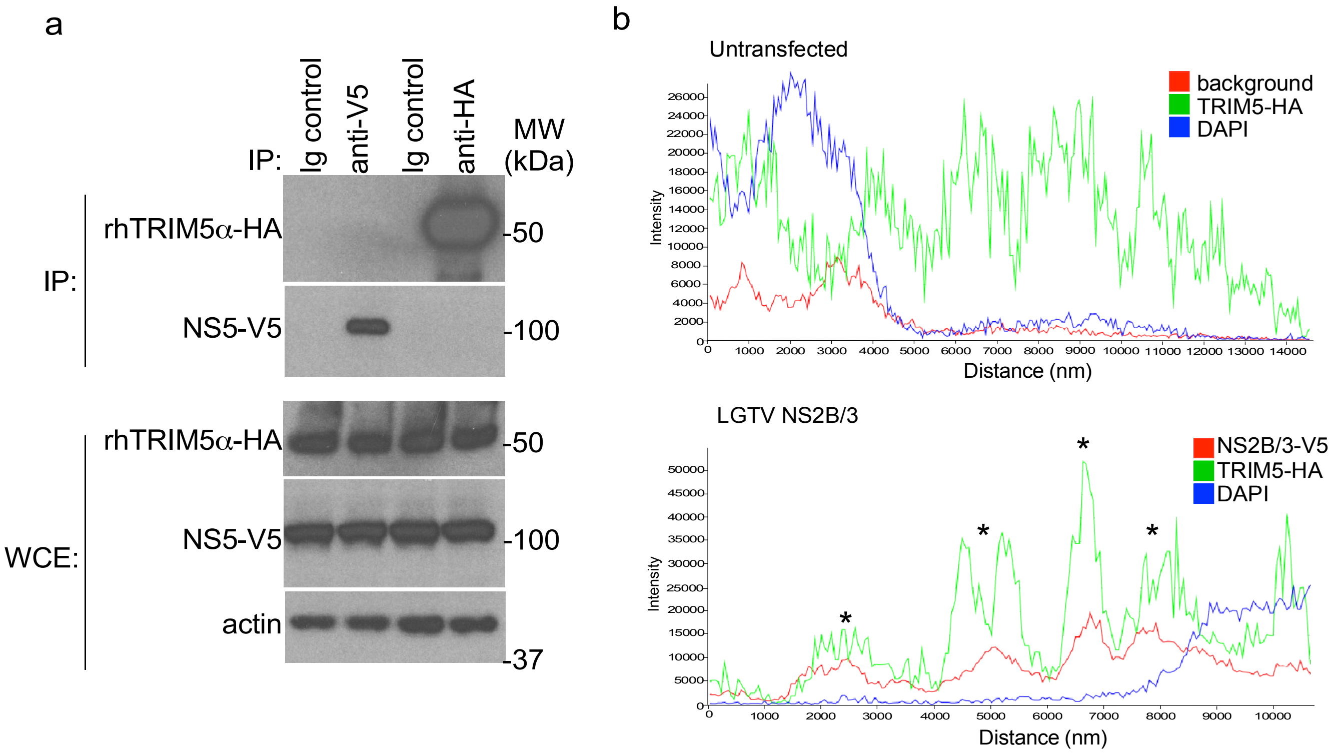
a
IP:
Ig control
anti-V5
Ig control
anti-HA
MW (kDa)
IP:
rhTRIM5α-HA
NS5-V5
-50
-100
WCE:
rhTRIM5α-HA
NS5-V5
actin
-50
-100
-37
b
Untransfected
background
TRIM5-HA
DAPI
Intensity
Distance (nm)
LGTV NS2B/3
NS2B/3-V5
TRIM5-HA
DAPI
Intensity
Distance (nm)

Suppl Fig 5

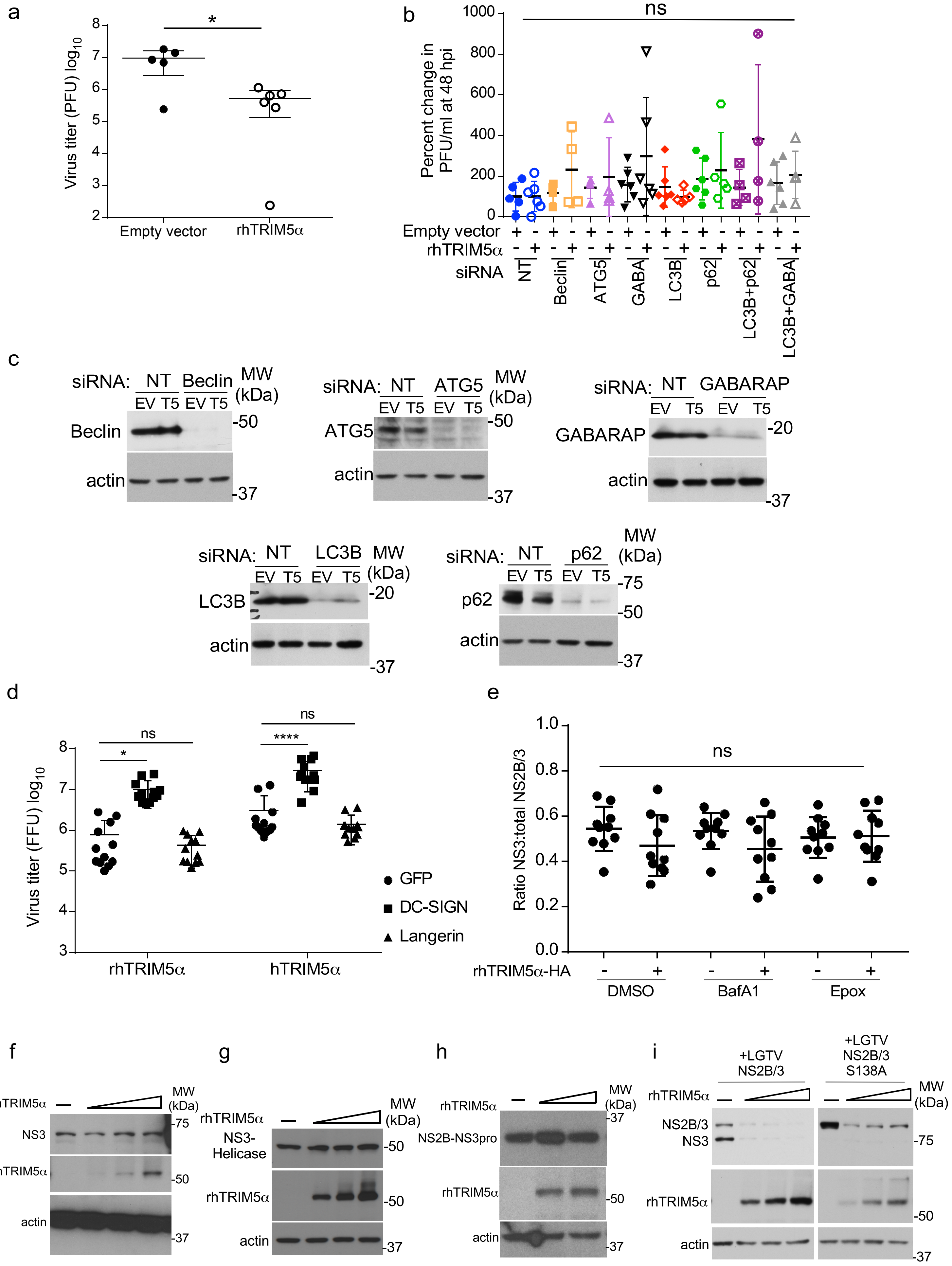

a
Virus titer (PFU) log10
Empty vector
rhTRIM5α
b
Percent change in PFU/ml at 48 hpi
ns
Empty vector
rhTRIM5α
siRNA
NT
Beclin
ATG5
GABA
LC3B
p62
LC3B+p62
LC3B+GABA
c
siRNA: NT Beclin
EV T5 EV T5
MW (kDa)
Beclin
actin
siRNA: NT ATG5
ATG5
siRNA: NT GABARAP
GABARAP
siRNA: NT LC3B
LC3B
siRNA: NT p62
p62
d
Virus titer (FFU) log10
GFP
DC-SIGN
Langerin
rhTRIM5α
hTRIM5α
e
Ratio NS3:total NS2B/3
rhTRIM5α-HA
DMSO
BafA1
Epox
f
rhTRIM5α
NS3
actin
g
NS3-Helicase
h
NS2B-NS3pro
i
+LGTV NS2B/3
+LGTV NS2B/3 S138A
NS2B/3
NS3

Suppl. Fig. 6

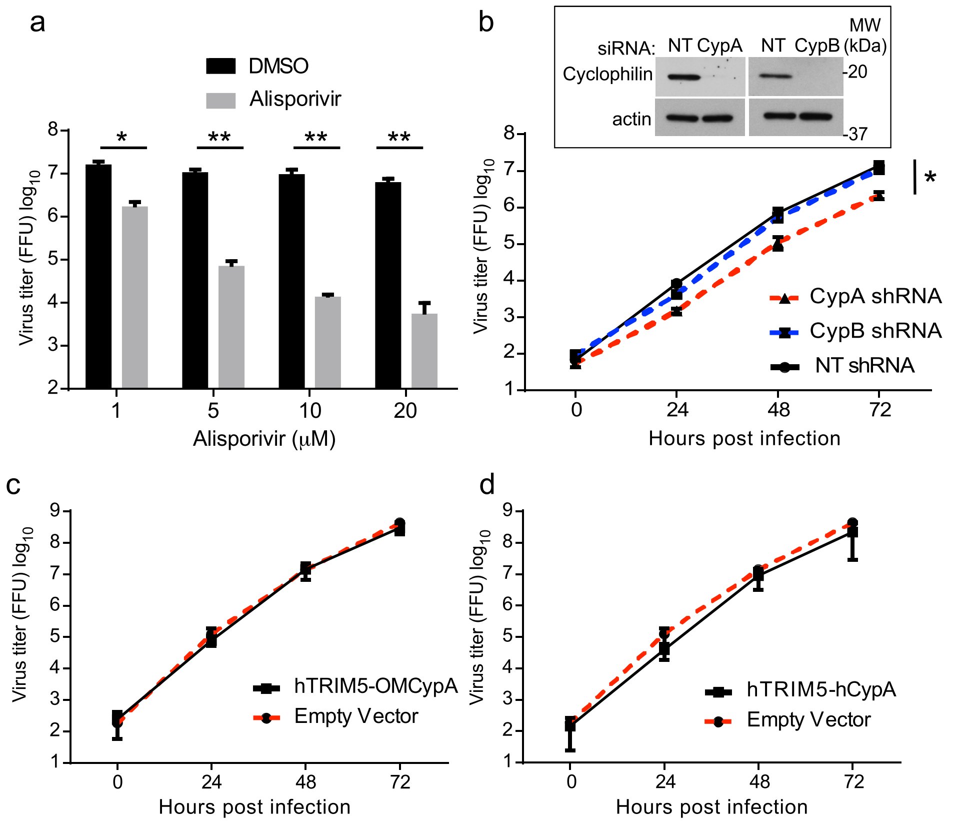
a
DMSO
Alisporivir
Virus titer (FFU) log10
*
**
**
**
1
5
10
20
Alisporivir (μM)
b
siRNA:
NT
CypA
NT
CypB
MW (kDa)
Cyclophilin
actin
-20
-37
Virus titer (FFU) log10
*
CypA shRNA
CypB shRNA
NT shRNA
Hours post infection
c
Virus titer (FFU) log10
hTRIM5-OMCypA
Empty Vector
Hours post infection
d
Virus titer (FFU) log10
hTRIM5-hCypA
Empty Vector
Hours post infection
